# Systematic characterization of site-specific proline hydroxylation using hydrophilic interaction chromatography and mass spectrometry

**DOI:** 10.1101/2023.07.28.550951

**Authors:** Hao Jiang, Jimena Druker, James W. Wilson, Dalila Bensaddek, Jason R. Swedlow, Sonia Rocha, Angus I. Lamond

## Abstract

We have developed a robust workflow for the identification of proline hydroxylation sites in proteins, using a combination of hydrophilic interaction chromatography (HILIC) enrichment and high-resolution nano-Liquid Chromatography-Mass Spectrometry (LC-MS) together with refining and filtering parameters during data analysis. Using this approach, we have combined data from cell lines treated with either the prolyl hydroxylase (PHD) inhibitor, Roxadustat (FG-4592), or with the proteasome inhibitor MG-132, or with a DMSO control, to identify a total of 4,993 and 3,247 proline hydroxylation sites, respectively, in HEK293 and RCC4 cells. A subset of 1,954 (HEK293) and 1,253 (RCC4) non-collagen proline hydroxylation sites with high confidence were inhibited by FG-4592 treatment. Features characteristic of proline hydroxylated peptides were identified, which were consistent between the HEK293 and RCC4 datasets. The more hydrophilic HILIC fractions were enriched in peptides containing hydroxylated proline residues, which showed characteristic differences in charge and mass distribution, as compared with either unmodified, or oxidised peptides. Furthermore, we discovered that the intensity of the diagnostic hydroxyproline immonium ion was dependent upon parameters including the MS collision energy setting, parent peptide concentration and the sequence of amino acids adjacent to the modified proline. We show using synthetic peptides that a combination of retention time in LC and optimised MS parameter settings allows reliable identification of proline hydroxylation sites in peptides, even when multiple prolines residues are present. Focussing on proteins in which newly identified proline hydroxylation sites were inhibited by the pan-PHD inhibitor, FG-4592, showed enrichment for proteins involved in metabolism of RNA, mRNA splicing and cell cycle regulation, including the protein phosphatase 1 regulatory subunit, Repo-Man (CDCA2). We show that Repo-Man is hydroxylated at P604 and in the accompanying study by Druker et al. ^1^, we present a combination of cellular and biochemical evidence that hydroxylation of Repo-Man at P604 is important for its function in controlling mitotic progression.

## Introduction

Site-specific hydroxylation of proline residues is an important enzymatic protein post-translational modification (PTM), which is mediated by prolyl hydroxylase (PHD) enzymes. For example, PHDs are essential for controlling HIF-α protein levels via their proline hydroxylation activity, in the presence of iron, 2-oxoglutarate and molecular oxygen ^2–4^. HIFs are key transcription factors that regulate the acute and adaptive responses to changes in oxygen availability through a variety of target genes involved in metabolism, proliferation, cell motility and cell death ^5–8^. Proline hydroxylation in HIF-α leads to an increased affinity with the VHL E3-ligase complex, K48-ubiquitination and proteasomal degradation ^9–11^.

Until recently, the HIF-α family was regarded as the only verified target for the PHD family of enzymes. Cummins et al ^12^ suggested that PHDs could also regulate the kinase activity of IKK enzymes, based upon functional assays in cells, but did not provide a formal demonstration mapping sites of proline hydroxylation in the IKK complex. More recently, several studies have reported other potential protein targets for PHDs ^12–20^, based upon either biochemical, and/or in-cell functional analyses, including mutations, and, in some cases, also mass spectrometry-based proteomics analysis ^21,22^. The use of MS-based proteomics analysis is widely established for the identification and characterisation of multiple PTMs, including phosphorylation ^23–25^, methylation ^26^, ubiquitination ^27^, glycosylation ^28,29^ and others ^30–33^. The benefit of using MS is that many modification sites can be identified and quantified in a robust and unbiased way^31^. However, so far it has proven difficult to reliably identify sites of proline modification in proteins, outside the collagen family, using MS methods. This technical challenge has contributed to a degree of controversy in the field regarding whether there are any intracellular targets for PHD-mediated hydroxylation beyond the HIF family of proteins. For example, a study by Cockman et al ^34^ reported that purified PHD enzymes failed to recapitulate proline hydroxylation *in vitro* at sites previously identified using mass spectrometry on proteins isolated from cells. They proposed, therefore, that HIF-α may be the only physiologically relevant target for PHDs. However, an alternative interpretation for their observations could be that technical limitations with the *in vitro* assays may be responsible for the failure to detect hydroxylation of non-HIF target proteins using purified PHDs.

The controversy surrounding whether there exist in cells *bona fide* PHD target proteins outside of the HIF family highlights several technical problems that complicate the identification of sites of enzymatic proline hydroxylation by MS. These include, (i) the lack of validated methods for enriching proline-hydroxylated peptides during sample preparation for MS and (ii) the risk of misidentification of hydroxylated proline, due to potential confusion with oxidation of Methionine (M), which can occur as a non-enzymatic, chemical modification of peptides during sample preparation. Other technical issues include possible confusion of hydroxylated proline with Leucine/Isoleucine (L/I) residues, (which have similar mass to Pro-OH), the frequent absence in spectra of fragment ions diagnostic for Pro-OH and other problems associated with low-quality MS spectra.

In previous studies using an MS-based proteomics workflow to identify proline hydroxylation sites, antibodies that recognise hydroxyproline were used to enrich either whole proteins, or peptides, prior to MS analysis ^35,36^, similar to antibody-based strategies commonly used to enrich other types of protein modifications for MS-based analysis. However, both the unreliability of currently available anti-hydroxyproline antibodies and the relatively low abundance of proline hydroxylated peptides at steady state, has, in practice, limited the success of this approach.

The hydroxylation of proline, which adds a hydroxyl (OH) group to the gamma carbon atom, significantly alters its chemical properties (Figure 1A). The pKa value in free proline, (Pro or P), is 10.8, while for hydroxylated Proline, (HyPro or P-OH), it is 9.68^37^. Hence, following proline hydroxylation, the resulting modified peptides will be more hydrophilic in acidic solution, which can be utilized for liquid chromatography (LC)-based separation.

**Figure 1:**
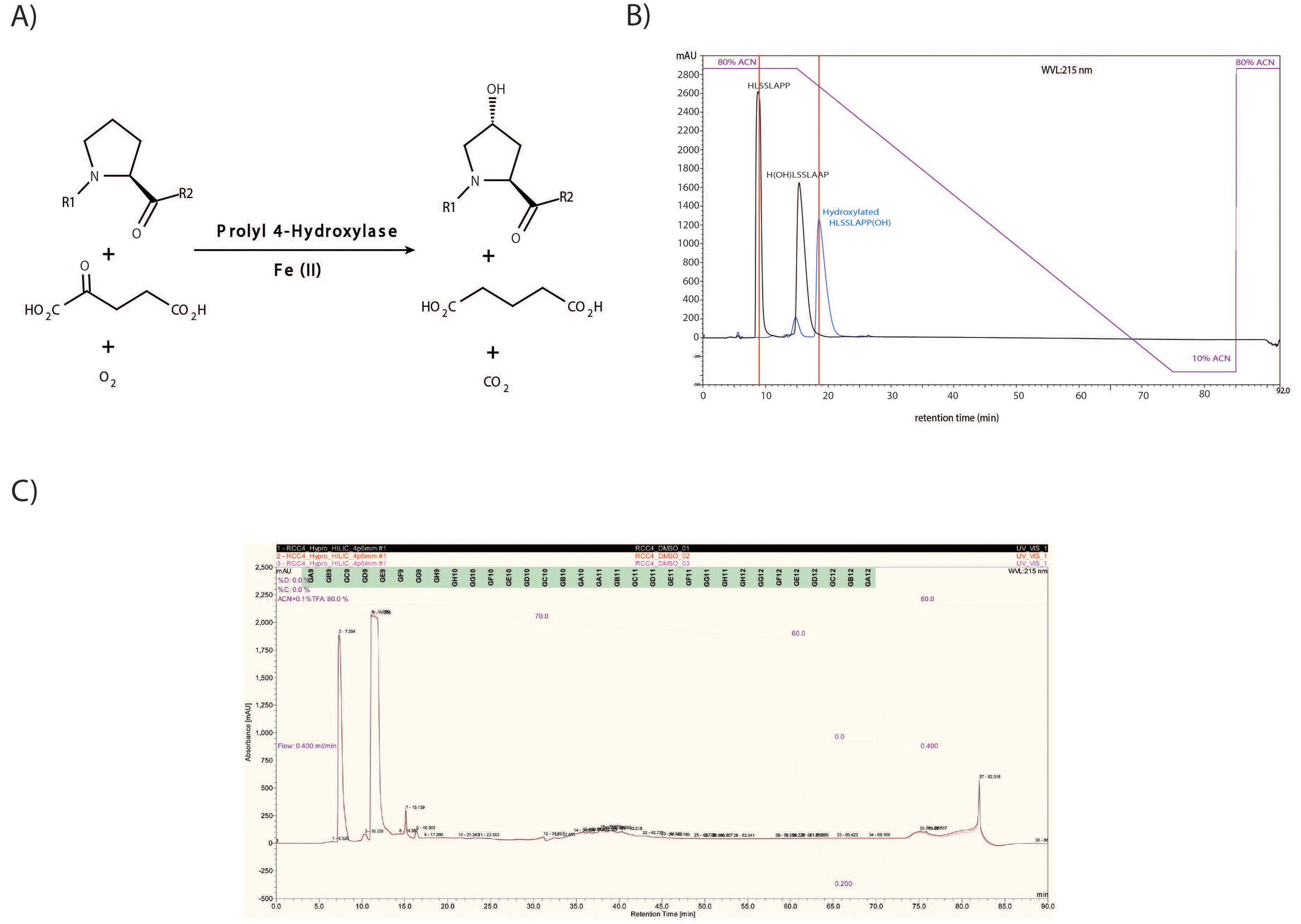
A) The schematic diagram of proline hydroxylation. B) HILIC chromatography showing the retention time difference of the synthetic peptides (HLSSLAPP, with no modification, or hydroxylation at the histidine/proline). C) HILIC chromatography profile of peptides fractionation from RCC4 samples (3 replicates, overlay).

In this study, we report the development of a robust approach for the enrichment and reliable identification of proline hydroxylation sites in proteins using an MS workflow that incorporates a chromatographic peptide enrichment step, using HILIC. This takes advantage of the increase in hydrophilicity that results from the addition of the hydroxyl group to proline. HILIC is known to be capable of separating polar compounds ^38^, including peptides ^39,40^ and carbohydrates ^41^ and has been reported previously to allow enrichment of hydroxylated peptides ^42^. We show that by combining HILIC enrichment with optimised MS instrument parameters and data analysis procedures, sites of proline hydroxylation can be reliably identified and subsequently verified by the analysis of synthetic peptides. Using this approach, we report the identification of 4,993 proline hydroxylation sites in HEK293 cell extracts and 3,247 sites in RCC4 cell extracts, along with analysis of common motifs found at sites of modification and classes of target proteins harbouring hydroxylated proteins. In the accompanying study by Druker et al. ^1^, we present functional data showing that the newly identified hydroxylation of Repo-Man (CDCA2) at proline 604 is important for its role in the control of mitotic progression, supporting a wider role for PHD enzymes in regulating cell cycle and potentially other cell functions.

## Results

We have previously compared chromatography methods for fractionating peptides with post translational modifications, prior to MS analysis, which showed that HILIC performed better than hSAX for enrichment of peptides containing hydroxylated proline residues ^42^.To extend this analysis, we first tested the performance of HILIC in separating synthetic peptides of identical length and sequence that were either unmodified, hydroxylated on histidine, or hydroxylated on proline. As shown in Figure 1B, proline hydroxylation increased peptide retention time on the HILIC column and thus proline hydroxylated peptides can be separated by HILIC from their corresponding unmodified form, while peptides with hydroxylation at different amino acids can also be separated.

The HILIC fractionation method was then applied to the analysis of tryptic peptide samples, generated from extracts of both RCC4 and HEK293 cell lines. Extracts were compared from cells treated with either the prolyl hydroxylase inhibitor, FG-4592, or with the proteasome inhibitor MG-132, to prevent degradation of proline hydroxylated proteins, with DMSO treated cell extracts used as a control. The chromatography profile, (Figure 1C, shadow green), demonstrates the reproducibility of HILIC fractionation. In total, 32 peptide fractions were collected, dried and directly used for LC-MS/MS analysis.

Using this HILIC enrichment strategy, by combining the results from the DMSO, FG-4592 and MG-132 datasets, a total of 3,588 and 5,491 distinct proline hydroxylation sites, (Table S1), were initially identified from the RCC4 and HEK293 samples, respectively. Next, by considering several unique features of HyPro, we established specific steps during data analysis, distinct from methods typically used for other PTM analysis, to enhance accurate identification of HyPro sites.

### HILIC fractionation distinguished proline hydroxylated peptides from oxidised peptides

MS analysis software usually identifies post-translational peptide modifications through the detection of the mass shift of a peptide and its corresponding fragment ions as a result of the modification. However, the hydroxylation of a proline equates to a mass shift of 15.994915 Da, which is similar to the mass shift seen for the oxidation of several other amino acids. For example, the oxidation of methionine residues is especially pertinent as it represents the amino acid on which chemical oxidation occurs most frequently. Indeed, methionine oxidation can either happen endogenously in cells, as a protection mechanism from oxidative damage on other reactive residues critical to the function of the protein ^43,44^, or it can occur exogenously, during sample preparation for MS analysis ^45^. Hence, one of the most controversial issues around the identification of proline hydroxylation sites is the possibility of confusing chemical oxidation on methionine, or other amino acid residues, for proline hydroxylation ^34^.

As shown above using synthetic peptides, HILIC chromatography has the potential to separate the peaks of proline hydroxylated peptides from others. We therefore compared the percentage/distribution of peptides identified as either unmodified, methionine oxidised, or proline hydroxylated, across 32 HILIC fractions derived from both HEK293 (Figure 2) and RCC4 (Figure S1) cell extracts. From the initial results of HEK293 samples, most unmodified peptides were identified in fractions F1-F20 (Figure 2A), with oxidised peptides showing a similar pattern, but more enriched in fractions F4-F14. In contrast, hydroxylated peptides were enriched in HILIC fractions F13-F19, consistent with their increased hydrophilicity, with the addition of a hydroxyl group causing increased retention time on HILIC. A similar HILIC fraction distribution pattern was also seen with the samples from RCC4 cell extracts (Figure S1A).

**Figure 2:**
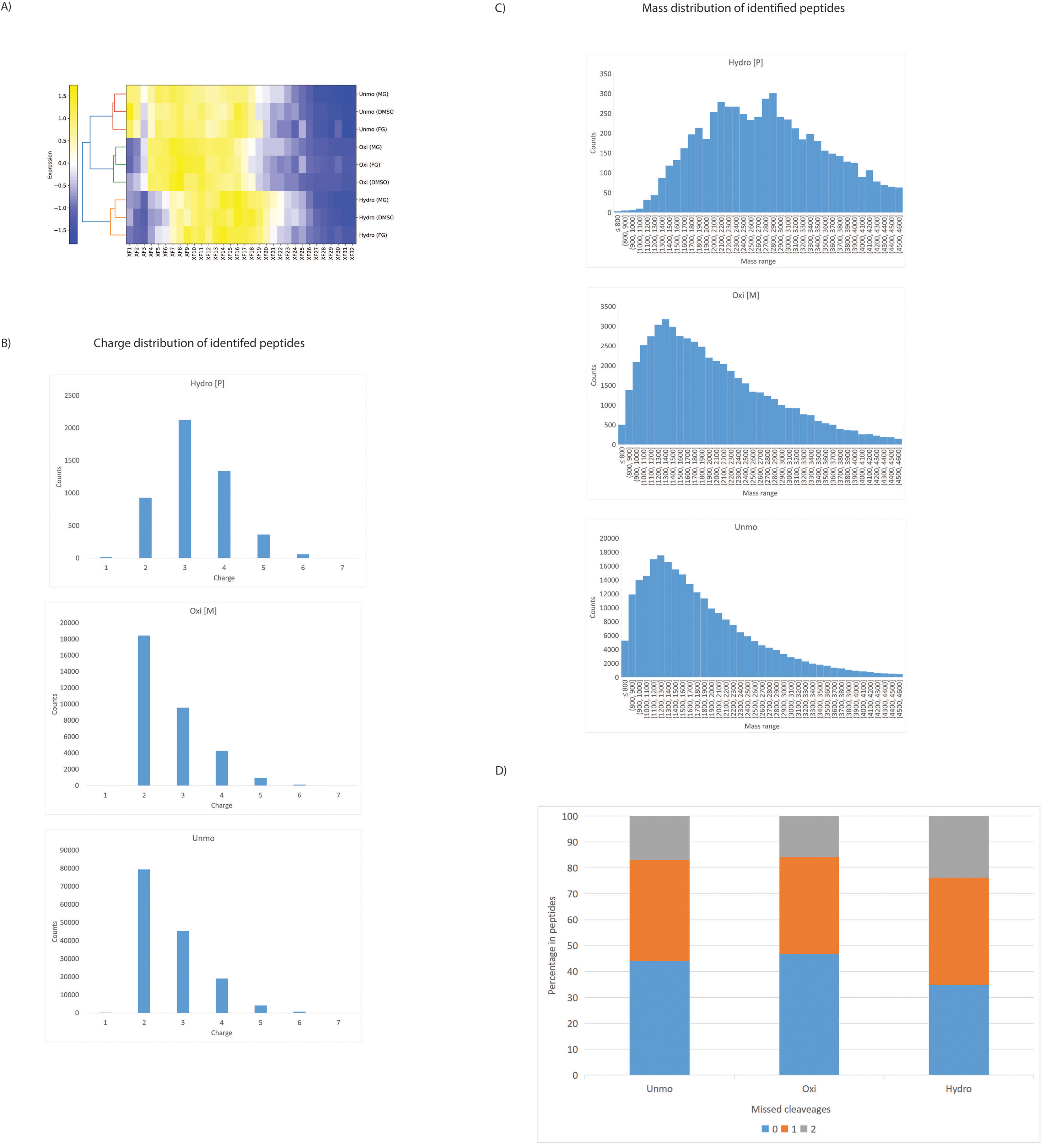
HEK293 dataset, A) The hydrophilicity difference of peptides with hydroxylation (P), oxidation(M) and unmodified peptides across HILIC fractions from all experiments with different treatment. To generate the heatmap, “modificationSpecificpeptides.txt” result file was used and divided into three files for identified peptides with either hydroxylation (P), oxidation(M) or unmodified peptides. The value of “Fraction x” column from of each peptide was used and summarized for each fraction, which represented the total quantity of peptides in each fraction. In each condition, the summarised fraction value of F1, F2, …F24 was scaled to 100% in total. The scaled fraction value were used for heatmap analysis by Heatmapper2 (Nucleic Acids Res. 2025 May 05. doi:10.1093/nar/gkaf385). The hydrophilicity increases from the Fraction 1 to Fraction 32 (XF1…XF32). B) The charge distribution of peptides with hydroxylation (P), oxidation (M) and unmodified peptides. C) The mass distribution of peptides with hydroxylation (P), oxidation (M) and unmodified peptides. D) The missed cleavage difference of peptides with hydroxylation (P), oxidation (M) and unmodified peptides.

Furthermore, the data also showed that the hydroxylated peptides have characteristic features that differ from either unmodified, or oxidised peptides. For example, we observed clear differences in both charge states, (Figure 2B and Figure S1B), and mass distributions, (Figure 2C and Figure S1C), of peptides containing hydroxylated proline residues. Thus, the most frequent charge state for both unmodified and oxidised peptide ions was 2+, with fewer peptide ions identified at higher charge states. In contrast, the most frequent charge state for peptide ions containing hydroxylated proline was 3+, with fewer peptide ions identified with a charge of either 2+, or 4+. The mass range for most of the unmodified and oxidised peptides was 800-2,220 Da, while for hydroxylated peptides it was 1,900-3100 Da.

A possible explanation for this observed variation in peptide mass value is that proline hydroxylation increases the frequency of missed cleavages when digesting proteins to peptides using trypsin, resulting in longer peptide sequences, with on average a correspondingly higher charge state and higher mass. Consistent with this hypothesis, a higher fraction, (∼65%), of the hydroxylated peptides from HEK293 samples were detected with at least one missed cleavage (Figure 2D), compared with the fraction of either unmodified peptides (56%), or oxidised peptides (53%). The same phenomenon was observed for RCC4 samples (Figure S1D), where 52% of the proline hydroxylated peptides were detected with missed cleavages, which was higher than the level of peptides for either unmodified peptides (48%), or oxidised peptides (38%).

In summary, HILIC fractionation is shown to provide a convenient approach for enriching hydroxylated peptides and separating these from both unmodified and oxidised forms of the same peptide sequences.

### Detection of HyPro diagnostic ion

The unique immonium ion (m/z of 86.0659), which is generated from the fragmentation of HyPro in MS, is known to distinguish HyPro from other amino acids in peptides, or PTMs (Figure S2) and thus can be diagnostic for the presence of a hydroxylated proline. However, we were intrigued to see that even for synthetic peptides that are known to include a HyPro residue, this immonium ion was not always detectable in the MS2 fragmentation spectra (Figure 3). We therefore investigated whether the instrument settings used for MS analysis might affect the generation and detection of the immonium ion, following fragmentation of different peptide sequences. To test this, we compared MS2 spectra obtained from the hydroxylated synthetic peptides LDLEMLAP(564)YIPMDDD; (derived from protein HIF1α ^3^), WHLSSLAP(1717)PYVK; (derived from protein CEP192 ^21^) and LAPITSDP(408)TEATAVGAVEASFK (derived from protein PKM2 ^14^).

**Figure 3:**
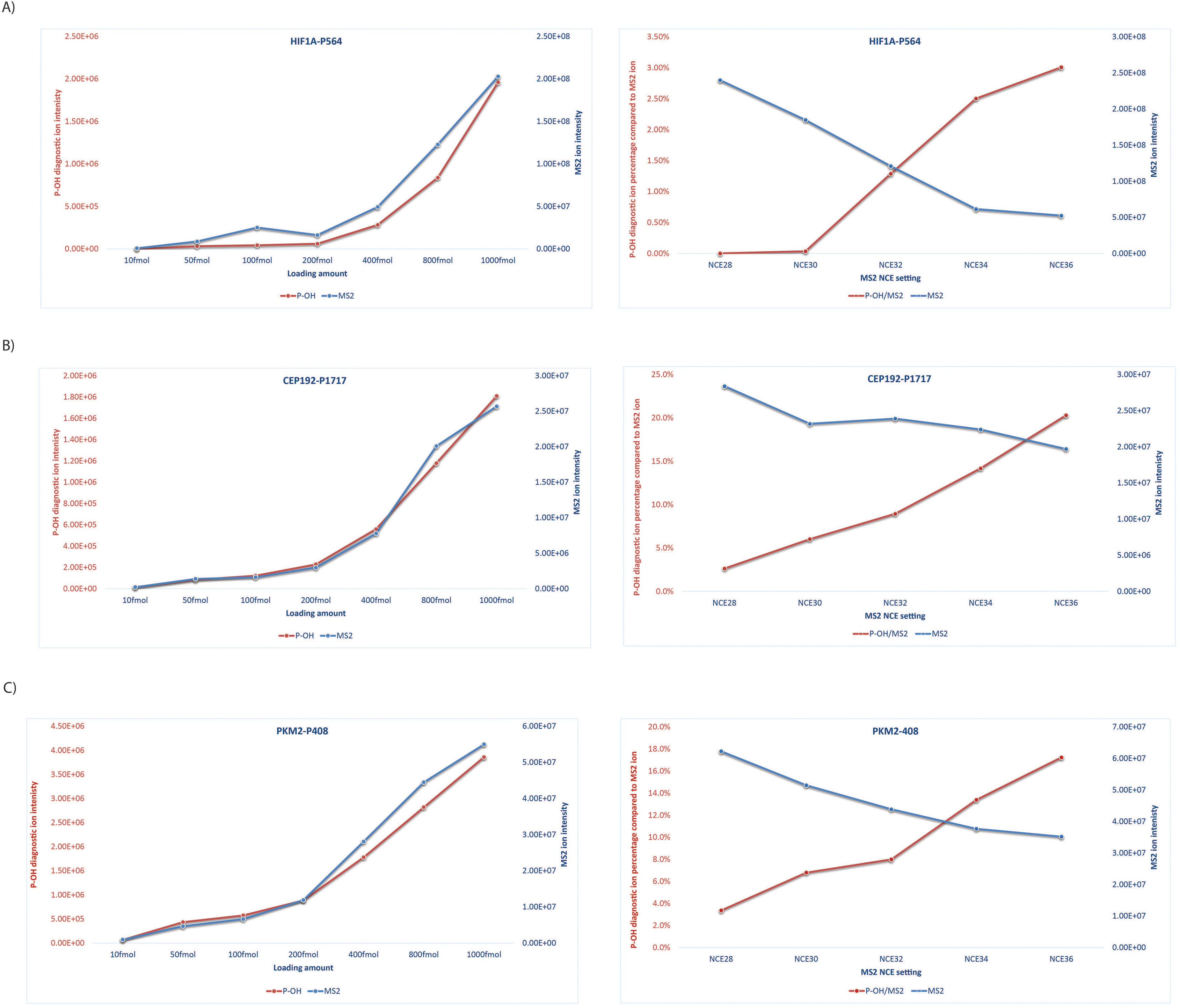
A) The intensity behaviour of HyPro diagnostic ion (m/z at 86.060) under different loading amount (left) and NCE settings (right), using the synthetic peptide HIF1α-P564, LDLEMLAPYIPMDDD (m/z at 833.9018). B) The intensity behaviour of HyPro diagnostic ion (m/z at 86.060) under different loading amount (left) and NCE settings (right), using the synthetic peptide CEP192-1717, WHLSSLAPPYVK (m/z at 707.38). C) The intensity behaviour of HyPro diagnostic ion (m/z at 86.060) under different loading amount (left) and NCE settings (right), using the synthetic peptide LAPITSDP (8) TEATAVGAVEASFK (m/z at 707.38). All ions were extracted under ±10ppm mass tolerance.

The intensity of related fragment ions was compared with each of these peptides (Figure 3). For all three peptides, (with 1 pmol of each analysed), the intensity of the diagnostic HyPro immonium ion, (relative percentage to base peak MS2 intensity), increased following an increase in the collision energy (NCE) setting in the MS instrument used to generate fragment ions from peptides, (Figure 3A, 3B and 3C), while a common NCE setting for HCD is ∼ 30 in an Orbitrap Tribrid MS platform.

Importantly, these results reveal that a higher than usual collision energy must be used during MS analysis to increase the probability of detecting the diagnostic HyPro immonium ion. We propose that this, along with other MS analysis parameters, (see below), may have contributed to some of the recent controversy in the field surrounding the ability to confirm the identification of PHD-dependent HyPro residues in non-HIF family protein targets.

Using a higher energy NCE setting increased the probability of generating the diagnostic HyPro immonium ion, while the low/medium energy settings generated more useful b/y ions that can used for peptide matching. However, even with equal loading and identical MS fragment settings, HyPro residues located within the context of different surrounding amino acid sequences showed variable intensity of the diagnostic HyPro immonium ion. Thus, for both the hydroxylated CEP192 peptide and PKM2 peptide, we readily observe ∼15-20% intensity of the HyPro immonium ion in MS2 spectra, while under identical MS analysis conditions the HIF peptide only generated ∼3% of the immonium ion at its maximum. We conclude that different peptides present different ionisation and transmission features in MS, depending on their amino acid composition and sequence, which in turn affects the ability to detect sites with HyPro modification.

Our studies have also identified other factors during MS analysis that can contribute to the efficiency of detecting the diagnostic HyPro immonium ion, including the concentration of the target peptide in the samples injected into the MS. If the target peptide amount is relatively low, even with optimal NCE settings, the number of diagnostic immonium ions transferred to the MS detector might fall below the detection limit. To test this hypothesis, different amounts of prolyl hydroxylated synthetic peptides were loaded and analysed by MS, using the target acquisition mode PRM: 10 fmol, 50 fmol, 100 fmol, 200 fmol, 400 fmol, 800 fmol and 1,000 fmol (Figure 3). As expected, both the MS2 base peak intensity and the diagnostic ion intensity, dropped as the loading amount on LC-MS decreased. Even with the more sensitive PRM mode, no diagnostic HyPro immonium ion was detected from the HIF1α peptide at 10 fmol (Figure 3A).

In summary, we have demonstrated that the intensity of the diagnostic HyPro immonium ion is sensitive to multiple factors involved in MS analysis, including collision energy settings in the MS instrument, the amino acid sequence surrounding the HyPro site in peptides and the quantity of the target modified peptide in the sample under analysis.

### Proline hydroxylation sites profiling of HEK293 and RCC4 cell lines

To further improve our workflow and reduce the chance of misidentification of HyPro, the MS data were filtered by the detection of the diagnostic HyPro immonium ion. For other potential HyPro sites, where the presence of the diagnostic immonium ion could not be confirmed, we recommend that a score cut-off and site localisation control should be used for identification. Accordingly, a score cut-off of 40 for hydroxylated peptides and a localisation probability cut-off of more than 0.5 for hydroxylated peptides was performed.

HEK293 (4 bio-replicates) and RCC4 (3 bio-replicates) cell lines were plated 24 h before treatment with either FG-4592 (50 µM, 24 h), or MG-132 (20 µM, 3 h before harvesting), or DMSO control (24 h). Cells were harvested, lysed, and further prepared for HILIC enrichment and LC-MS/MS analysis.

Implementation of the above parameters resulted in the identification of 4,768 and 3,063 proline hydroxylation sites without the associated diagnostic ions, and 4,993 and 3,247 sites in total, (together with the sites with diagnostic ions), from HEK293 and RCC4 samples, respectively (Table S2). These are considered as high confidence identifications.

Within the high confidence dataset, we successfully identified known proline hydroxylation sites in both cell lines, including the multiple proline sites of collagen proteins COL2A1 and COL4A1 and P62 in the ribosomal protein RPS2. Also, previously reported potential PHD-dependent hydroxylated proline residues were identified in either the HEK293, or RCC4 datasets, such as P294 of NDRG3 ^46^ , P319 of PPP2R2A ^47^ and P307/P322 of ACTB ^48^.

By following the diagnostic ion filter, 225 and 184 HyPro sites were confirmed from all the HEK293 and RCC4 samples, respectively (Table S2). Interestingly, more proline hydroxylation sites were found from collagen proteins in the RCC4 dataset (116 of 184), as compared with the HEK293 dataset (53 of 225). The hydroxylated peptide sequence in the collagen proteins shared the collagen-like sequence motif GxPGxx with Gxx repeats (Figure 4A and 5A). These data are not surprising, since HyPro is known to be a common PTM found in collagen proteins and critical for helix stability ^49,50^. No consistent sequence motifs were identified in the remaining 68 (HEK293) and 172 (RCC4) sites from non-collagen proteins (Figure 4A and 5A), which may indicate that these proteins include multiple distinct target classes, rather than all belonging to the same class of targets.

**Figure 4:**
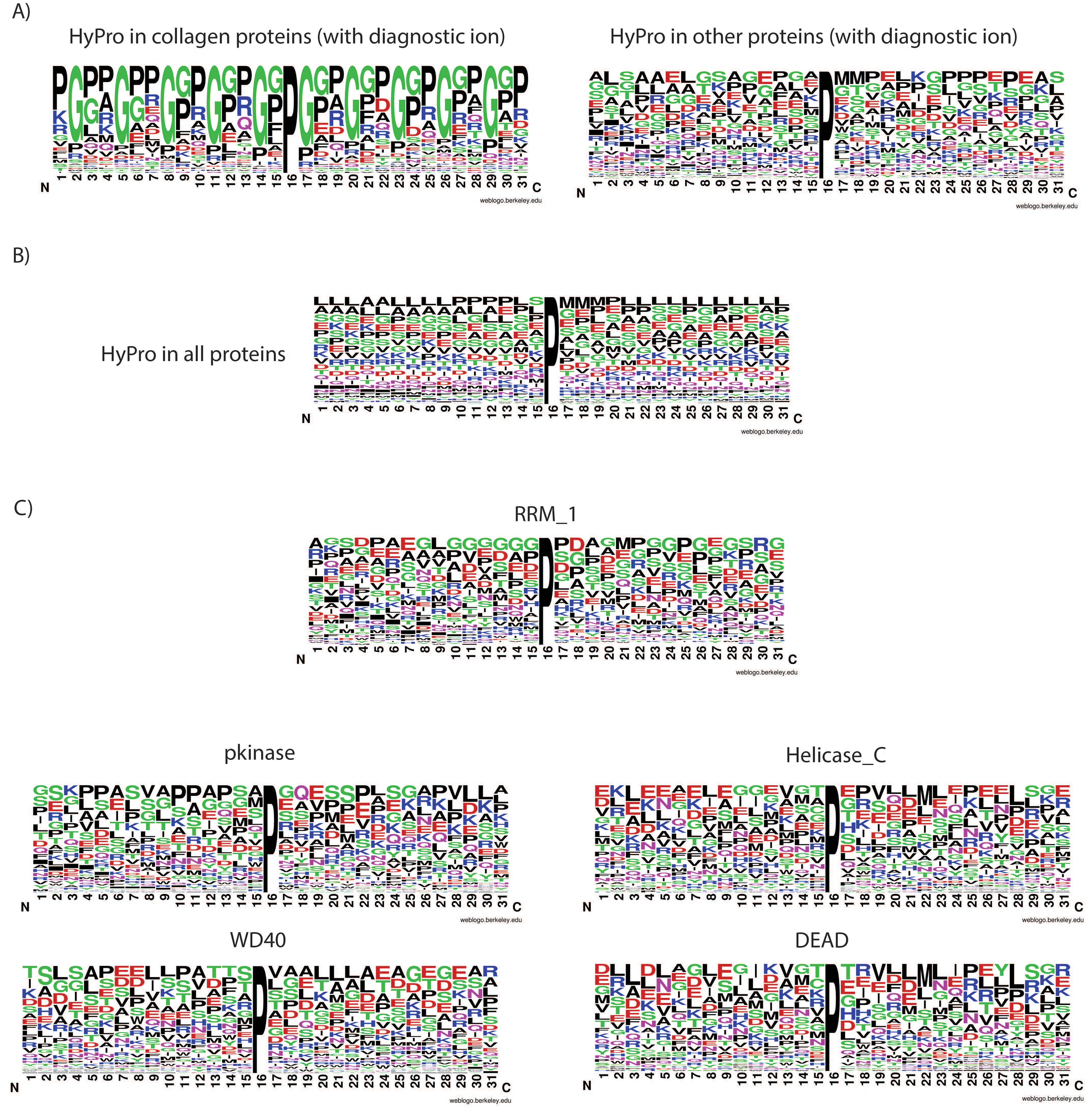
A) Sequence Logo for peptide window with hydroxylated proline site from collagen proteins (with diagnostic ion) and from other proteins (with diagnostic ion). B) Sequence Logo for peptide window with hydroxylated proline site from all the proteins. C) Sequence Logo for proline hydroxylated peptides from proteins included in different protein families (RRM_1 domain, pkinase domain, Helicase_C domain, WD40 domain and DEAD domain).

**Figure 5:**
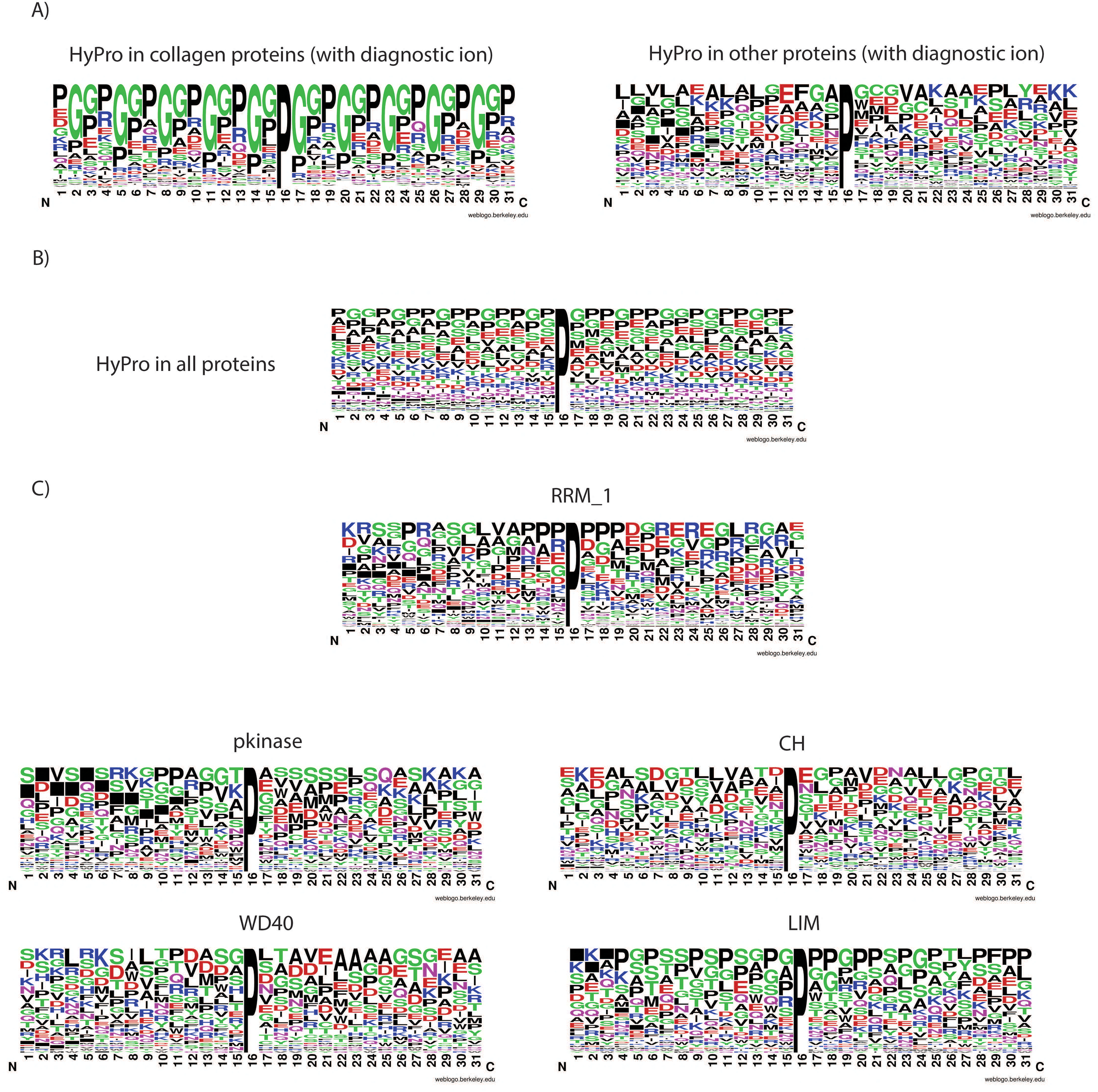
A) Sequence Logo for peptide window with hydroxylated proline site from collagen proteins (with diagnostic ion) and from other proteins (with diagnostic ion). B) Sequence Logo for peptide window with hydroxylated proline site from all the proteins. C) Sequence Logo for proline hydroxylated peptides from proteins included in different protein families (RRM_1 domain, pkinase domain, CH domain, WD40 domain and LIM domain).

Furthermore, to show the representation of the sequence conservation of amino acids in all the proline hydroxylated peptides sequence window, sequence logo analysis was performed (Figure 4B and 5B). A high frequency of the amino acids L, P, G, S and E in the hydroxylated peptides was immediately apparent in the HEK293 dataset. The RCC4 dataset showed a similar pattern, but without L. These data show that two different cell lines show unique profiles of proteins with hydroxylated peptides. This is an additional argument against these identifications of hydroxylated prolines arising via non-specific chemical oxidation during sample preparation, since an identical sample preparation procedure was used for both cell extracts.

We noted that there was a high frequency of a methionine residues appearing either at the first, second, or even third positions after the HyPro site (Figure S3). Bearing in mind concerns that HyPro identification by MS could be confused with oxidation of methionine, (see above), since both involve a mass shift of +16 Da, as a conservative approach these peptides were filtered out and a set of 3,521 and 2,728 filtered HyPro sites (Table S3), was generated for sequence analysis for the HEK293 and RCC4 datasets, respectively.

Next, we matched the proline hydroxylated proteins against the Pfam database (https://pfam-legacy.xfam.org/^51^), to investigate protein families and the potential presence of any overrepresented sequence motif. After the removal of all the collagen proteins, the most common terms identified in both samples included the RNA recognition motif (RRM_1), WD40 repeat, conserved site (WD40) and protein kinase domain (Pkinase). Besides, the DEAD/DEAH box helicase domain (DEAD) and Helicase, C-terminal (Helicase_C) were also observed in the HEK293 dataset, while the Calponin homology domain (CH) and LIM domain (LIM) were more enriched in RCC4 dataset. As shown in Figures 4C and 5C, there was no evidence for the over-representation of a single, simple motif in any of the protein families, but we noted that the amino acid frequency differed for different protein families. In the HEK293 dataset, the amino acids G, S, D and E were frequently located in proximity to the hydroxylated P residue for the hydroxylated peptides from RRM_1, while more L and A residues were in the hydroxylated peptides from WD40. For the Helicase_C domain, E was noticed frequently in the hydroxylated peptides. In the RCC4 dataset, multiple P, G and S amino acids were included in the sequence for the LIM domain, akin to collagen proteins, where proline hydroxylated peptides typically have multiple P and G. Also, a GTP motif was present in the proline hydroxylated peptides for the Pkinase domain. A previous study^22^ has also reported the proline embedded in the L/xGxP consensus sequences of the kinase domains, (DYRK1A and DYRK1B), as the target of hydroxylation by PHD1. This is a highly conserved motif, present in most of the kinases comprising the CMGC family.

### Reactome pathway analysis of the FG-4592 dependent dataset

Next, to identify cellular functions and pathways in which proteins with PHD-catalysed proline hydroxylation may be involved, we compared HyPro sites identified in extracts prepared from both HEK293 and RCC4 cell lines that had been treated with either (i) the prolyl hydroxylase inhibitor FG-4592, or (ii) the proteasome inhibitor MG-132, with DMSO treatment as the negative control. Reactome pathway analysis was performed using the HyPro sites list, (Table S4), filtered by the following conditions: (i) HyPro sites only identified in extracts from either DMSO, or MG-132 treated cells, but not in extracts from cells treated with FG-4592; (ii) HyPro sites from all collagen proteins were removed. All the subsequent pathway analyses were performed using g:Profiler ^52^.

Interestingly, the data from HEK293 cell extracts revealed a statistical enrichment for proline hydroxylated proteins belonging to two main categories: (i) metabolism of RNA, i.e., processing of capped intron-containing Pre-mRNA and mRNA splicing; (ii) cell cycle regulatory pathways, i.e., cell cycle (mitotic), S Phase, M Phase, Mitotic Anaphase, Mitotic G1 phase and G1/S transition (Table S5 and Figure 6A). The equivalent data from the RCC4 cell line also revealed proteins enriched in the same metabolism of RNA and cell cycle (mitotic) pathways, (Table S5 and Figure 6B), but additionally included enrichment of proteins involved in signalling by Rho GTPases, Axon guidance and L1 family of cell adhesion molecules (L1CAMs) interactions ^53^. By comparing the HEK293 and RCC4 datasets, the most highly enriched pathways in common to both cell lines are metabolism of RNA, cell cycle (mitotic) and cellular response to heat stress/stress/stimuli.

**Figure 6:**
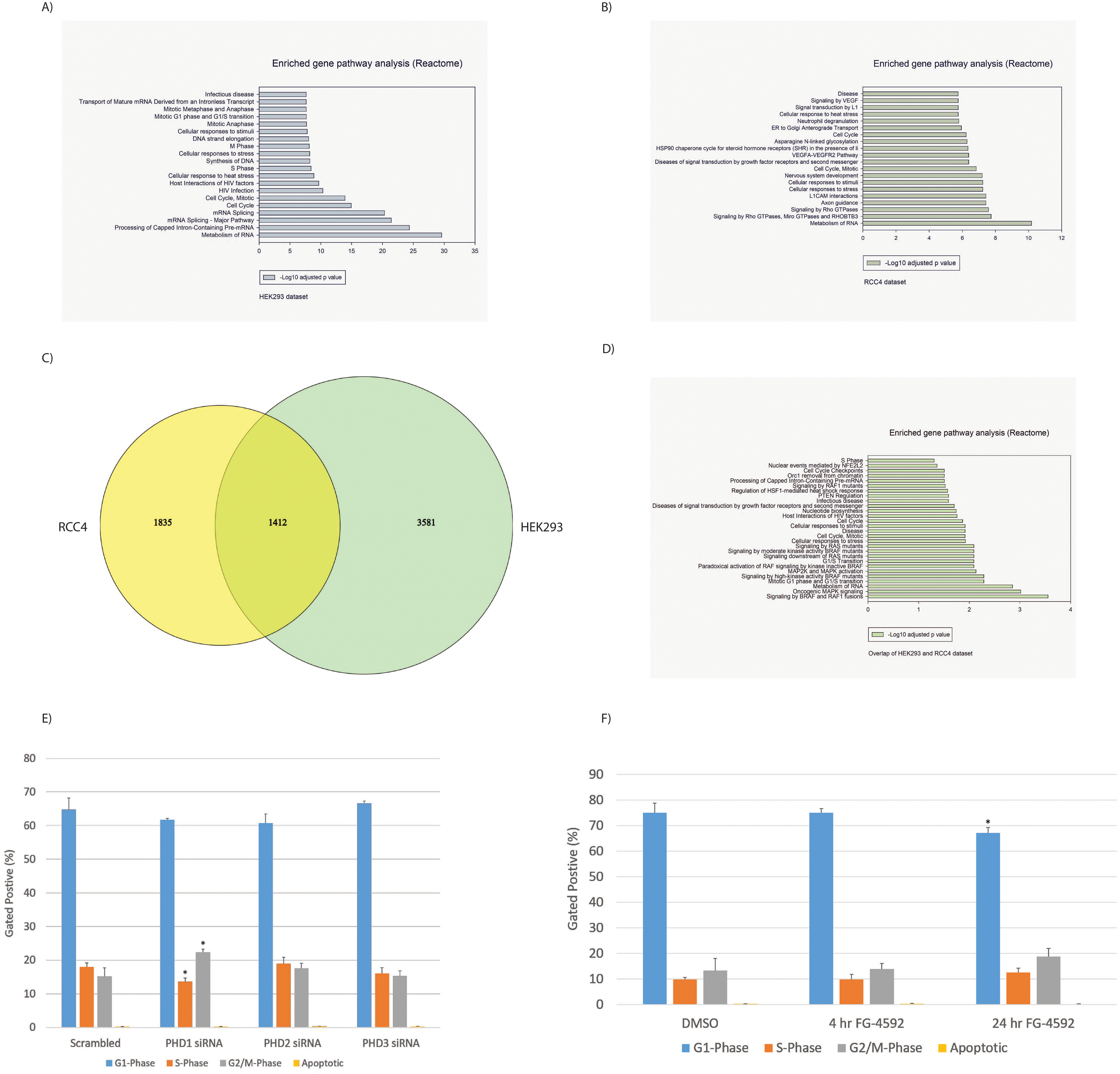
A) Statistical gene enrichment pathway analysis (Reactome) of the proteins only hydroxylated in DMSO/MG-132 treated HEK293 cells but not in FG-4592 treatment. B) Statistical gene enrichment pathway analysis (Reactome) of the proteins only hydroxylated in DMSO/MG-132 treated RCC4 cells but not in FG-4592 treatment. C) The Venn diagram of the identified proline hydroxylation sites between HEK293 and RCC4 dataset. D) Statistical gene enrichment pathway analysis (Reactome) of the proteins, which not only have the same hydroxylation proline sites from HEK293 and RCC4 dataset (DMSO/MG-132 treated), but also not hydroxylated in any FG-4592 treatment. E) RCC4 cells were treated with the indicated siRNAs and the cell cycle profile was determined by flow cytometry. The values presented represent the averages from three independent experiments. The error bars indicate the standard deviations. F) RCC4 cells were treated with the PHD inhibitor FG-4592 for the indicated times points prior to fixing. After which the cell cycle profile was determined by flow cytometry. The averages from three independent experiments are presented, whilst the error bars indicate the standard deviations. The cell profiles were gated according to the control in each independent experiment and the number of cells in each phase are expressed as a percentage of the total number of cells present. Statistical analysis was performed according to the Student’s T test vs the control; *p < 0.05.

In total, 1,412 identical proline hydroxylation sites were identified in both the HEK293 and RCC4 datasets, (Figure 6C, generated on http://barc.wi.mit.edu/tools/venn/). After filtering to remove sites at which hydroxylation was not inhibited by FG-4592 treatment, 167 proline hydroxylation sites, (corresponding to 136 proteins), were identified in both cell lines as strong candidates for being PHD targets (Table S4). By analysing Reactome pathway for the proteins in which these sites were located, (Table S5 and Figure 6D), metabolism of RNA and cell cycle regulatory pathways were enriched again. This is consistent with, a previous study ^21^ that has also shown that the proline hydroxylation of the centrosomal protein Cep192, mediated by PHD1, could be directly linked to the control of cell cycle progression.

RCC4 cells have a mutant VHL gene, where HIF1α and HIF2α cannot be targeted for proteasomal degradation, thus providing a model for analysis of VHL-independent mechanisms related to cell cycle progression. To determine whether PHD enzymes influence cell cycle progression, independent of VHL and HIF-α stabilisation, each of the three PHD isoforms were independently depleted by siRNA and analysed by fluorescence-activated cell sorting (FACS). Interestingly, following depletion of PHD1 in RCC4 cells, the G1-phase and S-phase populations decreased, while the G2/M populations significantly increased (Figure 6E). In contrast, the knockdown of PHD2 and PHD3 resulted in no significant change in the respective levels of the different cell cycle populations. Similarly, 24-hour treatment of RCC4 cells with the PHD inhibitor, FG-4592, resulted in a significant reduction in the G1-phase population and an increase in cells in the G2/M-phase, (Figure 6F), consistent with the PHD1 knockdown data.

### Validation of proline hydroxylation sites using synthetic peptides

While the above data provide robust identification of HyPro sites in proteins, even using HILIC fractionation enrichment and score/probability cut-off, it cannot be completely excluded that some false proline hydroxylation identifications. Therefore, we have included an extra validation step for identification of hydroxylated peptides using synthetic peptides to characterise cognate MS spectra for HyPro-containing peptide sequences. This focussed on the following criteria:

1. Comparison of the LC retention time of the experimentally measured peptides and corresponding synthetic peptides.
2. Investigation of MS1 and MS2 ion features, including manual analysis of the spectra to confirm the results obtained from software, determining the fragment b/y ions with high quality, and noting the ions neighbouring the hydroxylation sites.
3. Identification of the diagnostic HyPro immonium ion.

This approach is illustrated for the peptide sequence: KPLLSPIPELPEVP(600)EM(602)TP(604)SIP(607)SIRR, derived from the protein Repo-Man (CDCA2), which was first discovered as a chromatin-associated regulator of protein phosphatase one (PP1) ^54^ and identified here as a potential PHD target from the HEK293 dataset (Table S1 and Table S4). In this peptide, multiple prolines (P600/P604/P607) could potentially be hydroxylated, but none had a localisation probability high enough (>90%) to confirm unambiguously the actual HyPro site. Manual inspection of the MS/MS spectra indicated that amino acid residues M602, P604 or P607 could be either oxidised, or hydroxylated, each causing a +16 Da mass shift. Some of the fragment ions in the MS/MS spectra were missing that would have allowed us to distinguish between hydroxylation of either P604, or P607 residues, or oxidation at M602. This can be a common situation when analysing peptides isolated from cell or tissue extracts, especially when investigating low-abundance peptides/proteins.

To distinguish between these possibilities, we first investigated how either the hydroxylation of P, or the oxidation of M, would alter peptide chromatography in RPLC-MS analysis. This was achieved by comparing a set of synthetic peptides that were either hydroxylated, oxidised, or unmodified and determining their respective retention times (Figure 7A). This revealed that the peptides hydroxylated at either P604, or P607, had retention times that peaked at 76.54 minutes and 77.37 minutes, respectively. By contrast, the retention time of the unmodified peptide was 80.26 minutes, while that of the methionine oxidised peptide was 74.30 minutes. We conclude that the retention time measured by RPLC thus provides a discerning feature for HyPro modified peptides that can be used to validate protein hydroxylation sites and reduce the chance of misidentification of methionine oxidation as a potential HyPro site.

**Figure 7:**
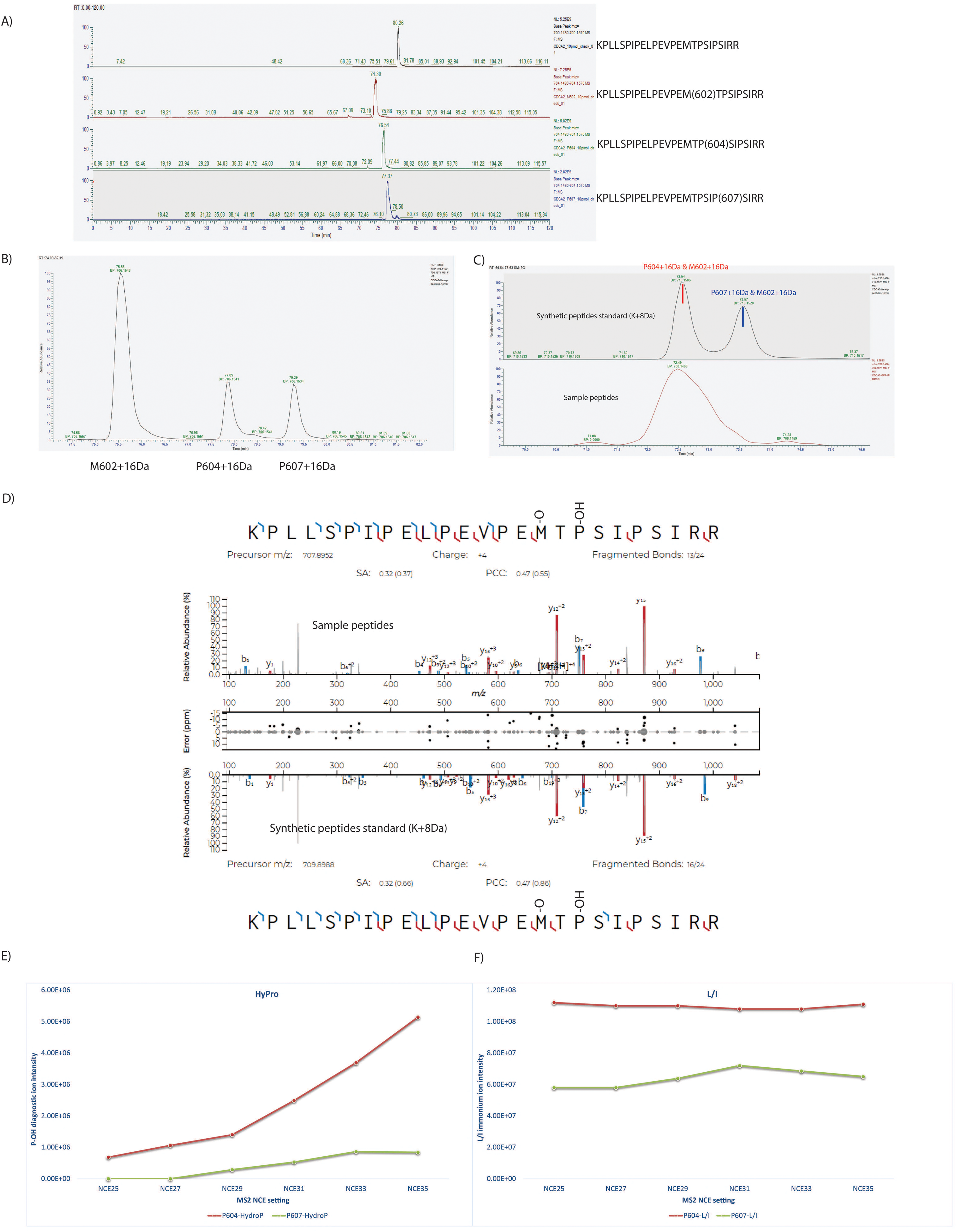
A) The retention time difference of the synthetic peptides (KPLLSPIPELPEVPEMTPSIPSIRR) with or without oxidation/hydroxylation (+16Da mass shift) at different amino acids using RPLC-MS analysis, individually. B) The retention time difference of the synthetic peptides (KPLLSPIPELPEVPEMTPSIPSIRR, K+8Da) with or without oxidation/hydroxylation (+16Da mass shift) at different amino acids using RPLC-MS analysis, mixed. C) The retention time comparison of the M602 oxidised peptides (KPLLSPIPELPEVPEM(O)TPSIPSIRR) with hydroxylation (+32Da mass shift) at different proline residues, between the spiked-in heavy isotope labelled synthetic peptide and the peptide in the IP sample. D) The MS/MS spectrum of the sequence KPLLSPIPELPEVPEM(O)TP(OH)SIPSIRR from the spiked-in heavy isotope labelled synthetic peptide (scan no. 21243) and the peptide from the IP sample (scan no. 25891). The intensity behaviour of E) HyPro diagnostic ion (m/z at 86.060) and F) L/I immonium ion (m/z at 86.097) under different NCE settings, using the synthetic hydroxylated peptide KPLLSPIPELPEVPEMTP(604)SIPSIRR and KPLLSPIPELPEVPEMTPSIP(607)SIRR (m/z at 704.15). All ions were extracted under ±10ppm mass tolerance.

To validate the MS1 and MS2 ion features of the prolyl-hydroxylated peptide, we next performed an IP-MS experiment in HeLa cells transiently transfected with a Repo-Man expression construct. For this analysis, the cells were arrested in Prometaphase by treatment with nocodazole. Cell extracts were prepared and the corresponding synthetic peptides, with heavy isotope labelled lysine, (K, +8Da), were spiked into each sample before LC-MS/MS analysis, to facilitate identification and quantification of the proline hydroxylated Repo-Man peptide. By measuring the RPLC retention time of synthetic peptides, the oxidation at M602 (75.55 min) and hydroxylation at either P604 (77.89 min), or P607 (79.29 min), could be distinguished, despite each having a similar mass shift of +16 Da (Figure 7B).

We note that the Repo-Man peptides hydroxylated at either P604, or P607, could be further modified by oxidation also at M602, causing a mass shift +32 Da (Figure 7C). By manually checking the experimental LC profile and MS/MS spectra (Figures 6C and 7D, generated via Universal Spectrum Explorer^55^), it was clear that a Repo-Man peptide could be detected with a mass shift of 32 Da, indicating that both hydroxylation at P604 and oxidation at M602 could occur at the same time. The site of modification was confirmed from the MS/MS spectra via comparison with the spectra from the synthetic peptides, based upon similarity in the respective MS2 fragmentation patterns (Figure 7D).

The diagnostic immonium ion was not detected from either of the proline hydroxylation site in these MS2 spectra, neither for the endogenous Repo-Man peptides from HeLa cells, nor for synthetic hydroxylated Repo-Man peptides. As described above, the detection of the HyPro immonium ion in MS spectra depends on the collision energy and the quantity of target peptides. Using the typical NCE setting of 27 on a Q Exactive MS platform, the diagnostic immonium ion of the synthetic Repo-Man proline hydroxylated peptide was barely detectable (Figure 7E). However, as the NCE setting was increased, there was a corresponding increase in the intensity of the HyPro diagnostic immonium ion detected. In contrast with the HyPro immonium ion, varying energy settings showed little to no effect on the efficiency of detection of the immonium ions from L/I (Figure 7F).

Finally, by comparing the RPLC retention times of the respective spiked-in heavy isotope-labelled synthetic peptides with hydroxylation at either P604 (72.54 min, with M602 oxidation), or hydroxylation at P607 (73.57 min, with M602 oxidation), these could be clearly distinguished. Therefore, we conclude that the Repo-Man protein in HeLa cell extracts is hydroxylated specifically at P604.

### Hydroxylation of Repo-Man at P604 is PHD-dependent

Next, we investigated further whether the hydroxylation of P604 on Repo-Man is PHD-dependent in cells. To this end we performed SILAC (Stable Isotope Labelling of amino acids in cell culture)-based quantitative proteomics in both HeLa and HEK293 cells, comparing cells grown +/− the PHD inhibitor FG-4592. Asynchronous cells were grown either in ‘heavy’ K4+R6 media, i.e., containing a combination of heavy isotope labelled lysine (K) and Arginine (R), or in ‘light’ K0+R0 (normal) media. Although HIF1α stabilisation in response to FG-4592 is efficient (Figure S4A), FG-4592 was added to cells growing in heavy media for 24h to ensure that the PHD activity was inhibited throughout the cell cycle, while DMSO was added in the light media as a control (Figure 8A). Total cell extracts were prepared, subjected to HILIC enrichment and the resulting samples were analysed by LC-MS/MS.

**Figure 8:**
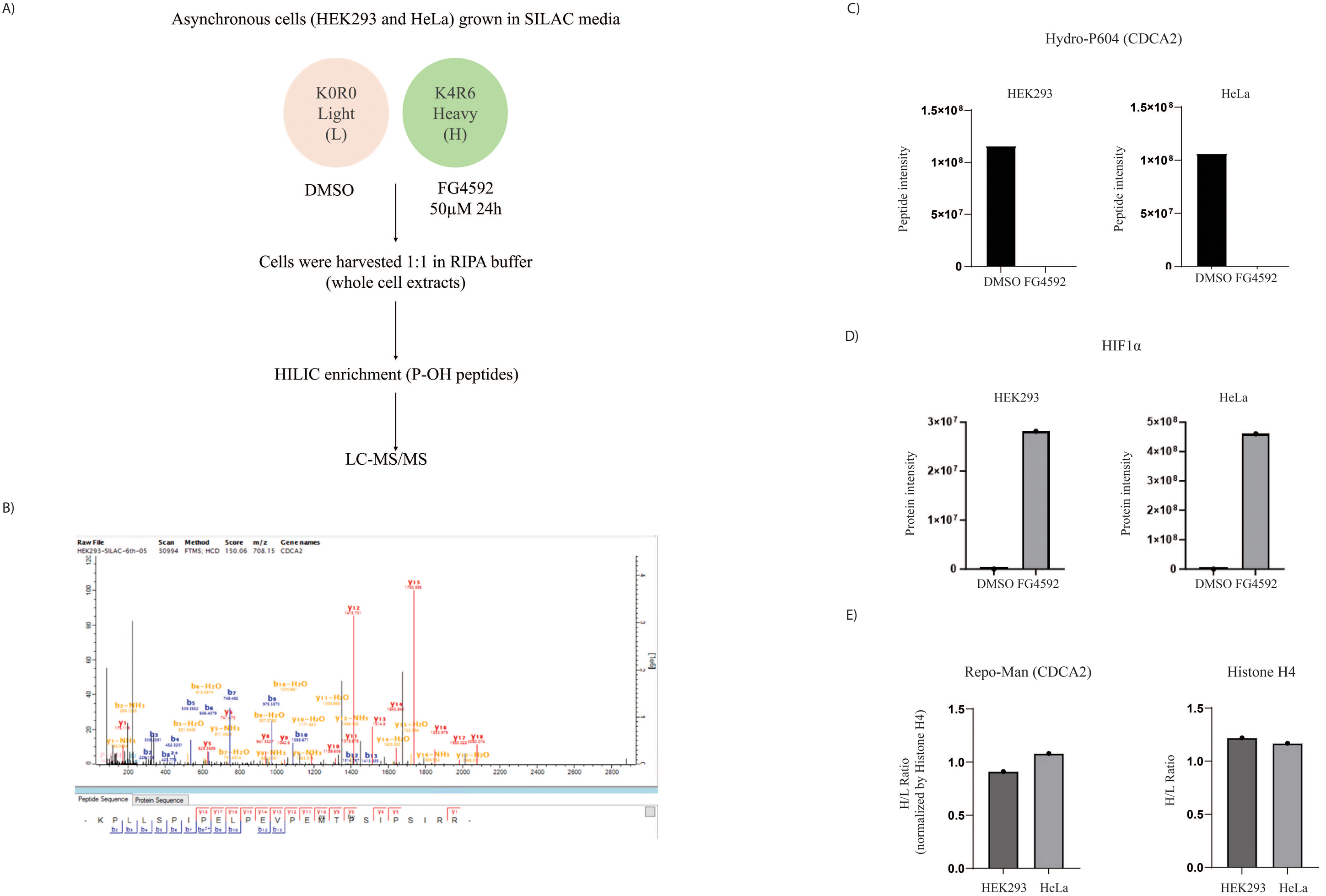
MS analysis of Repo-Man hydroxylation at P604. A) Diagram of proline hydroxylation sites analysis in asynchronous HeLa and HEK293 cells using SILAC, Cells were grown in SILAC media for 6 passages and 24 h before harvesting cells were treated with FG-4592 50 µM or DMSO. DMSO treated cells (control) were light labelled, and FG-4592 treated cells are heavy labelled. B) MS/MS spectra of endogenous Repo-Man peptide (KPLLSPIPELPEVPEMTPSIPSIRR), where matched fragment ions (b/y) were marked. Both the peptide precursor ion (m/z 708.15) and its corresponding fragment ions (y8 and y10) confirmed the identification of KPLLSPIPELPEVPEM(602-Ox)TP(604-OH)SIPSIRR. The diagnostic ion (m/z 86.06) of hydroxylated proline was also shown as “P-OH”. C) Quantification analysis of P604 hydroxylation in Repo-Man in both HeLa and HEK293 cells, under the treatment of either DMSO or FG-4592. The bar chart was based on the intensity of the peptide containing P604 hydroxylation. D) Quantification analysis of HIF1α in both HeLa and HEK293 cells, where the protein raw intensity was used for the bar chart since there’s no HIF1α detected in DMSO treated cells. E) Quantification analysis of Repo-Man (CDCA2) and Histone H4 in both HeLa and HEK293 cells, where H/L represents the heavy to light ratio using SILAC quantification.

The spectrum of the Repo-Man tryptic peptide (KPLLSPIPELPEVPEMTPSIPSIRR) containing hydroxylated P604, oxidised M602 and the diagnostic ion is shown in Figure 8B. The abundance of the hydroxylated (OH-P604) Repo-Man peptide in each condition i.e., DMSO and FG-4592, was quantified with its intensity shown in Figure 8C. The Repo-Man peptide hydroxylated at P604 was only detected in extracts of cells grown in the DMSO control media and not in extracts from cells grown in the presence of the PHD inhibitor FG-4592 (Figure 8C).

As a control for the FG-4592 treatment, we quantified levels of HIF1α, which, as expected, was only detected in the FG-4592 treated samples (Figure 8D). Furthermore, the FG-4592-dependent differences observed in Repo-Man hydroxylation in HeLa and HEK293 cells were not due to changes in total Repo-Man protein levels, as the ratio H/L is ∼ 1 (Figure 8E), confirming no significant protein level alterations in cells treated with FG-4592. As an additional control for the possible effect of SILAC media on protein expression, we measured Histone H4 levels, which demonstrated no significant changes in response to FG-4592 treatment (Figure 8E). In summary, these data show that Repo-Man hydroxylation at P604 in cells is PHD-dependent.

### Recombinant PHD1 hydroxylates Repo-Man at P604 *in vitro*

To determine whether Repo-Man could be a substrate for PHD hydroxylation *in vitro*, we tested if recombinant PHD1 would hydroxylate a synthetic Repo-Man peptide at the P604 site that was hydroxylated in cell extracts (Figure 9, A-B). To avoid any misinterpretation between oxidation and hydroxylation, we used a synthetic Repo-Man peptide as a substrate that was already oxidised at M602 (KPLLSPIPELPEVPEM(OX)TPSIPSIRR). Following incubation *in vitro* with recombinant PHD1, samples were then analysed by LC-MS/MS. Figure 9A compares the MS2 spectra obtained from incubating the Repo-Man peptide with recombinant PHD1, together with a reference synthetic peptide that is hydroxylated on P604. By comparing the fragment ions from MS2 spectra obtained from incubating the Repo-Man peptide +/− recombinant PHD1, especially the Y5 and Y8 fragment ions derived from the parent ion, (KPLLSPIPELPEVPEM(OX)TPSIPSIRR +16Da), we confirm that the PHD1 hydroxylation site is P604 and not P607 (Figure 9, A-B).

**Figure 9:**
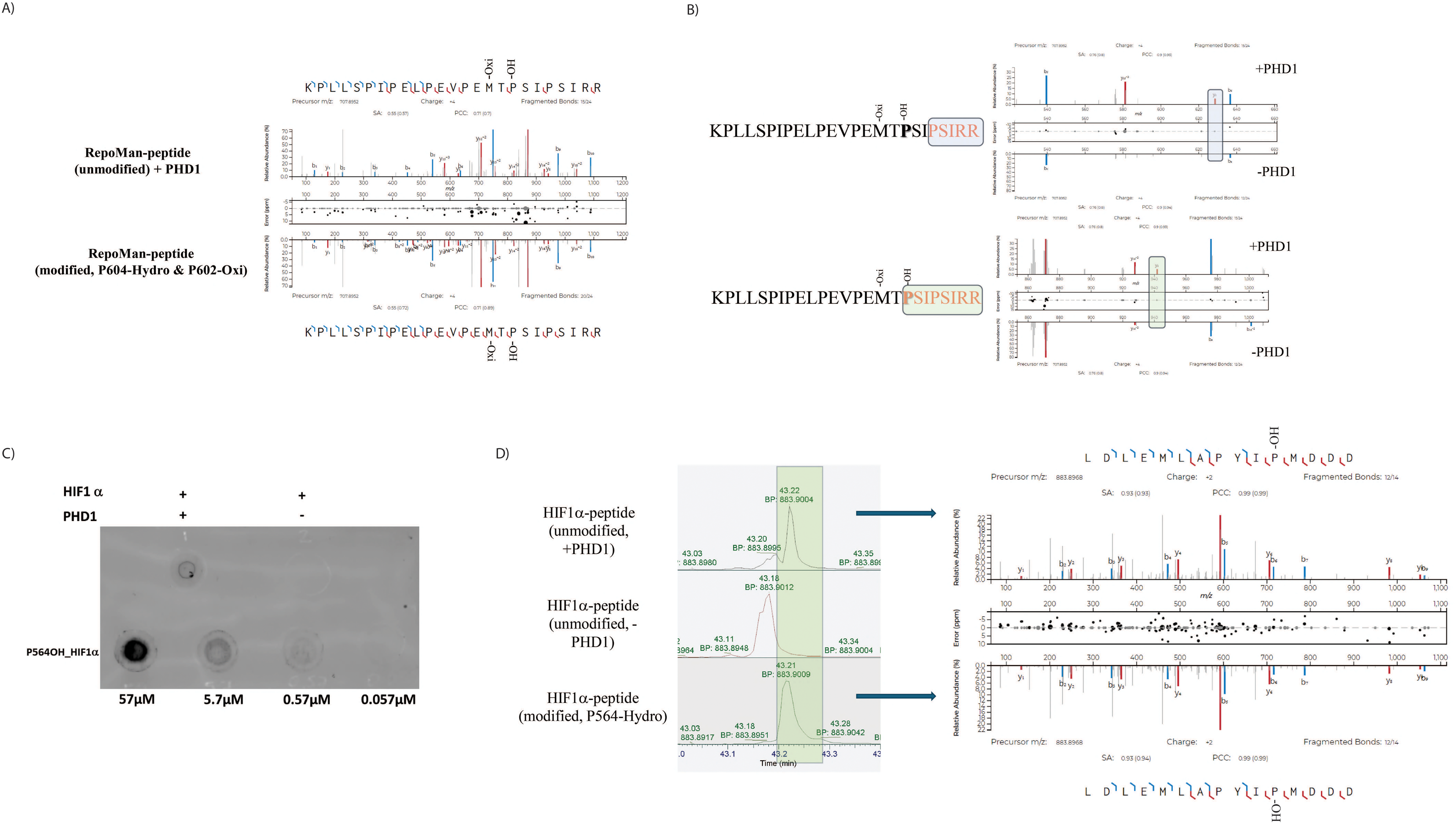
*In vitro* hydroxylation of Repo-Man synthetic peptide by recombinant PHD1. The reactions were performed in parallel using Repo-Man (KPLLSPIPELPEVPEM(602-OX)TPSIPSIRR) and HIF1α (LDLEMLAPYIPMDDD) synthetic peptides as substrates in combination with (+) or without (-) recombinant PHD1. A) MS/MS spectra of the hydroxylated peptide KPLLSPIPELPEVPEM(602-OX)TP(604-OH)SIPSIRR from Repo-Man. The mirror image shows the fragment ions matching results between the experimental hydroxylated peptide (top) and its synthetic standard peptide (bottom), where matched b/y ions were highlighted. B) Detailed MS/MS spectra of the hydroxylated synthetic peptide KPLLSPIPELPEVPEM(602-OX)TP(604-OH)SIPSIRR from Repo-Man. The mirror image shows the fragment ions matching results of the synthetic peptide incubated with or without PHD1. The y8 ion represents the fragment of [P(604-OH)SIP(607)SIRR], while the y5 ion represents the fragment of [P(607)SIRR], which confirmed the hydroxylation takes place at P604 only when incubation with PHD1. C) Dot blot analysis of the in vitro hydroxylation of HIF1α peptide, with and without recombinant PHD1 (upper dots). Titration of the OH-564 HIF1α peptide as a control of the primary antibody. Membrane was incubated with anti-OH-564 HIF1α and developed using far-red secondary antibody in typhoon. D) LC-MS analysis of *in vitro* hydroxylation of HIF1α peptide (LDLEMLAPYIPMDDD). The left panel shows the retention time difference in LC separation profile of HIF1α synthetic peptide with or without PHD1, compared to the HIF1α hydroxylated (P546-OH) synthetic peptide. The mirror image in the right panel shows the MS/MS fragment ions matching results between the experimental hydroxylated HIF1α peptide (top) and its synthetic standard peptide (bottom), where matched b/y ions were highlighted.

As a control, we performed in parallel the *in vitro* hydroxylation using the HIF1α peptide, (QDTDLEMLAPYIPMDDDFQLR) (Figure 9, C-D). Taking advantage of the availability of an antibody that recognises OH-P564 in HIF1α, a dot blot was performed with part of the *in vitro* reaction. As a control for the OH-564-HIF1α antibody, varying concentrations of the HIF1α synthetic peptide were tested on the dot blot (Figure 9C). The rest of the sample was analysed by MS. Figure 9D shows the MS spectra of HIF1α, with a +16Da shift only seen when the peptide was incubated with PHD1 and with the resulting hydroxylated peptide having a similar retention time to that observed for a positive control OH-564-HIF1α synthetic peptide. We note that the efficiency of the *in vitro* hydroxylation reaction, using only recombinant PHD1, was significantly higher with the HIF1α peptide than with the Repo-Man peptide. It is possible that the hydroxylation efficiency for Repo-Man *in cellulo* is increased by other factors, and/or by the sequence context of the endogenous protein, which are not reflected in the simplified *in vitro* assay.

In summary, the data demonstrate that a Repo-Man peptide can be *in vitro* hydroxylated at P604 by recombinant PHD1, consistent with data above showing that Repo-Man is a target for PHD hydroxylation in Hela and HEK293 cells.

## Discussion

In this study, we have successfully established a robust and reproducible workflow for the reliable identification of proline hydroxylation sites in proteins, using mass spectrometry analysis (Figure 10). This workflow includes a sample preparation step to enrich proline hydroxylated peptides, prior to LC-MS/MS analysis. Enrichment is achieved using HILIC fractionation, which exploits the change in hydrophilicity of peptides resulting from addition of a hydroxyl group on proline.

**Figure 10:**
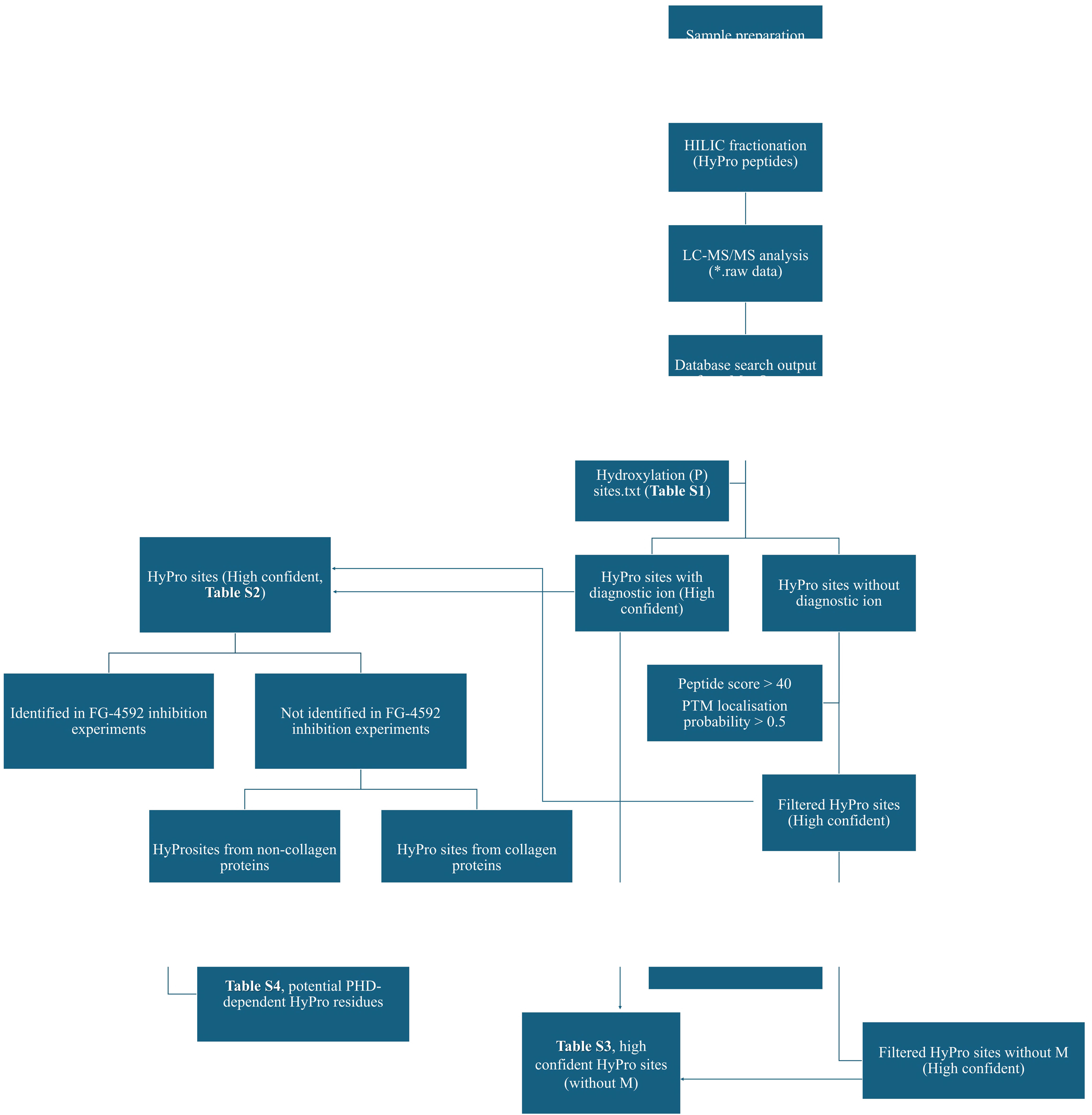
A flowchart summary of our workflow to identify proline hydroxylation sites.

In addition to hydroxylated peptide enrichment using HILIC fractionation, we have identified several parameters involved in the MS protocol and subsequent data analysis that affect the ability to detect specific proline hydroxylation sites and distinguish these from alternative forms of amino acid oxidation. For example, tryptic peptides modified with HyPro were found to show differences in peptide mass and charge state, as compared with either unmodified peptides, or peptides with oxidised methionine.

We note that it was previously reported to be difficult to distinguish between peptides with either HyPro (C_6_H_13_NO_2_), or Leu/Ile (C_5_H_9_NO_3_), because their corresponding residue mass difference is only 0.03639 Da (Figure S2). However, using modern MS instruments for proline hydroxylation identification, this concern is mitigated, as modern instruments are now capable of high mass resolution at both MS1 and MS2 levels. For example, using the Orbitrap Fusion MS, the MS1 and MS2 resolution values were 70,000 and 15,000, respectively. This corresponds to 10 ppm and 20 ppm as the mass tolerance parameter in data searching software and the most recent MS instruments provide even higher resolution. This provides sufficient mass resolution to distinguish whether a peptide contains either HyPro, or Leu/Ile.

Of particular interest, we have found that the generation of the diagnostic HyPro immonium ion in MS2 spectra is very sensitive to the collision energy setting used for peptide fragmentation and is also sensitive to the amino acid sequence of the peptide and its concentration in the analysis sample. In contrast, other immonium ions, e.g., for Leu/Ile, do not show this sensitivity to collision energy and are detected with similar efficiency over a broad range of energy settings. We propose that these observations can explain, at least in part, why the diagnostic HyPro immonium ion was only detected for 4.5% of the HyPro sites we mapped here from HEK293 cell extracts and 5.6% for RCC4 in the hydroxylation profiling experiment. This discovery, regarding the sensitivity of MS analysis parameters for generation and detection of diagnostic HyPro immonium ions, may be a significant contributory factor in some of the recent controversies surrounding the identification of proline hydroxylated PHD target proteins.

As shown in this study, the use of synthetic peptides, combined with optimised analysis procedures, provides a reliable strategy for identifying cellular proteins modified by proline hydroxylation. Thus, we recommend that for HyPro site mapping and for the purpose of validating specific target sites, a workflow is used that combines higher NCE energy settings in MS, (combined with optimisation using synthetic peptides), using a highly sensitive MS scan mode, (such as SIM, PRM for target analysis), to increase the detection limit and using HILIC enrichment to increase the concentration of HyPro modified peptides in samples being analysed.

We show that HILIC chromatography analysis of synthetic peptides can help to distinguish cases where proline hydroxylation could potentially be confused with oxidation of adjacent methionine residues. The similar mass shift resulting from proline hydroxylation and methionine oxidation is a technical challenge that can potentially lead to misidentifications in MS studies. Using synthetic peptides, specific proline hydroxylation sites in target proteins can be distinguished from methionine oxidation, based upon differential chromatographic behaviour of peptides with either hydroxylated proline or oxidised methionine, as well as by detailed analysis of fragmentation spectra. However, in the case of our global peptide analysis, as we were not able to perform synthetic peptide comparisons for every putative site identified, for comparative analyses we took the pragmatic approach of filtering out examples of peptides where a methionine residue was present within three residues of a potential proline hydroxylation site. This was done simply to reduce the possibility of misidentification in the set of novel proline hydroxylated peptides identified but as a consequence may also remove peptides that include bona fide proline hydroxylation sites.

Having validated the analysis strategy for detecting peptides modified with proline hydroxylation, we proceeded to characterise HyPro sites in extracts of both the HEK293 and RCC4 human cell lines. Several data analysis filters were applied to compare the sites, e.g., removing peptides mapped to collagen proteins, removing peptides containing methionine residues close to the HyPro sites, as explained above and removing peptides where the proline hydroxylation sites were still observed after treating cells with PHD inhibitors. Using these filtered peptides, sequence motif analysis showed that proline hydroxylation sites were enriched in the RNA recognition motif (RRM_1), protein kinase domain (Pkinase) and WD40 repeat, conserved site (WD40).

Given the potential wider biological importance of PHD-mediated regulation of key cellular processes, it is worth considering how the identification of proline hydroxylated target proteins can be further improved in future. For example, although we show here that proline hydroxylated peptides can be usefully enriched by HILIC fractionation, prior to LC-MS/MS analysis, we note that the enrichment efficiency is not as high as that possible with some methods for enrichment of peptides carrying other forms of PTM, particularly in the case of enrichment for phosphorylated peptides using TiO_2_ ^56^ or IMAC chromatography ^57^. HILIC fractionation will also enrich peptides with increased hydrophilicity due to modifications such as phosphorylation and glycosylation ^58^, which might complicate the detection of hydroxylated peptides. It will be interesting, therefore, to test whether the enzymatic removal of other modifications during sample preparation, such as using PNGase F to remove the N-glycans from proteins ^59^, might further affect the efficiency of proline hydroxylation identification using HILIC. Another issue worth noting is the high abundance of proline hydroxylated peptides derived from collagen proteins. Since MS is a concentration-sensitive analysis tool, high abundance ions from the collagen proteins may inhibit the ionisation of others, lower abundant proline hydroxylated ions derived from other, medium/low abundant proteins, thereby reducing the chance of their identification. To mitigate this problem, it may be helpful to include extra steps during sample preparation to selectively remove collagen proteins/peptides, prior to MS analysis. Furthermore, as previously mentioned, proline hydroxylated proteins tend to show increased levels of missed cleavages during trypsin digestion. A strategy of using multiple proteases, other than trypsin, has previously been used to increase the identification depth in proteomics studies ^60,61^ and this may also be interesting to test in future for identification of proline hydroxylation sites.

We show here that a Repo-Man peptide containing P604 and the surrounding sequence is recognised and hydroxylated *in vitro* by purified, recombinant PHD1, but note that the efficiency of this *in vitro* hydroxylation is lower than that seen for hydroxylation of a HIF1α peptide. Based upon the lower efficiency seen for *in vitro* hydroxylation of the RepoMan peptide by recombinant PHD1 alone, we suggest that *in cellulo* the activity of PHD enzymes towards many target proteins may depend upon additional targeting subunits. This would be analogous to the role of targeting subunits for protein phosphatase PP1 and PP2 catalytic subunits ^62–65^, such as Repo-Man itself and B56 ^66,67^. The efficiency of hydroxylation may also be increased by other features not replicated in the basic *in vitro* system, e.g. post-translational modifications of residues on the target sequence surrounding the hydroxylation site. In this regard it is interesting that even for the canonical PHD target HIF1α, the LIMD1 protein was identified as a bridging factor between PHDs and HIF1α ^68^. It will be interesting in future to investigate this further with experiments to evaluate factors that can stimulate the efficiency of site-specific hydroxylation *in vitro* on Repo-Man and other PHD targets. Reactome pathway analysis showed that the proteins identified as targets of proline hydroxylation in both the HEK293 and RCC4 cell lines were enriched preferentially for factors involved in cell cycle regulation and RNA processing, suggesting possible roles for PHD-mediated regulation in both these processes. This is consistent with our previous studies showing that the regulation of cell cycle progression can be controlled by mechanisms involving PHD-mediated proline hydroxylation, including HyPro modification of the centrosomal protein CEP192 ^21^. With regard to cell cycle regulation, it is interesting that we also detected in this study hydroxylation of P604 in the mitotic regulator Repo-Man, which is a chromatin-associated subunit of a protein phosphatase one complex. We show further that hydroxylation of Repo-Man at P604 is PHD-dependent in cells and that this site is a substrate for hydroxylation by recombinant PHD1 *in vitro*.

In the accompanying study by Druker et al. ^1^, we provide detailed experimental evidence showing that hydroxylation of Repo-Man at P604 is of functional importance for controlling progression through *mitosis*, acting, at least in part, via hydroxylation of Repo-Man at P604 regulating the interaction of Repo-Man with the B56-PP2A phosphatase complex during chromosome alignment and thereby controlling the levels of Histone H3T3 phosphorylation. In our opinion, these data, showing the functional importance of the novel PHD-catalysed proline hydroxylation site at P604 on Repo-Man, together with the large number of additional proline hydroxylation sites in multiple proteins identified in this study, are not consistent with the recent suggestion made by Cockman et al., who proposed that HIF-α may be the only physiologically relevant target for PHDs. This argument was based, at least in part, on the finding that HIF-α is a much better substrate for efficient hydroxylation by recombinant PHD1 than other targets, in a highly purified *in vitro* assay system. As discussed above, an alternative explanation is that many physiological substrates for hydroxylation by PHDs may require additional subunits and/or co-factors for efficient hydroxylation to be recapitulated *in vitro*.

Based upon the many novel proline hydroxylation sites we have identified in multiple target proteins, we propose that future experimental studies may show further important roles for PHD-mediated cellular regulation linked with stress responses and oxygen sensing, particularly relating to the control of RNA processing and cell cycle progression.

## Materials and Methods

### Cell culture

HEK293, RCC4 and HeLa cells were cultured in Dulbecco’s Modified Eagle Medium (Gibco, # 41966-029), supplemented with 10% Fetal Bovine Serum (FBS, Gibco # A3169801), 100 U/ml penicillin and streptomycin and 2 mM L-Glutamine. Cell lines were maintained at 37℃ with 5% CO2 in humidified incubator.

HEK293 or RCC4 cells were plated in 10-cm plates and cultured for 24 h. DMSO or FG-4592 (Selleck # S1007, 50 µM) was then added into the media for the following 24 h incubation. MG-132 (Calbiochem, 20 µM) was added for 3 h.

For SILAC ^69^, HEK293 and HeLa cells were cultured in DMEM for SILAC media (Thermo scientific # 88364), supplemented with 10% Dialyzed FBS (Gibco #A33820), 100 U/ml penicillin, streptomycin, 2 mM L-Glutamine and either ‘”Light” amino acids (K0R0, Lysine and Arginine), or “Heavy” amino acids (K4R6, ^13^C_4_-Lysine and ^13^C_6_-Arginine). The media was changed every 1–2 days. Cells were kept in SILAC media for 6 cell passages to achieve >95% labelling. Before the last passage, cells were plated in 10-cm plates and 24 h later. Either DMSO or 50 µM FG-4592 (Selleck # S1007), were added for the last 24 h in light, or heavy media, respectively. After incubation, cells were washed with PBS and trypsinised. Then cells from each condition were resuspended in PBS. Cells from both conditions were mixed at 1:1 ratio (cell number) and centrifuged at 1,000 g to collect the pellets.

### Lysates of HEK293, RCC4 and HeLa cells

HEK293, RCC4 and HeLa cells were harvested in PBS and the pellets were lysed in 500 µL of lysis buffer (50 mM Tris pH 7.5, 150 mM NaCl, 1% NP40, 0.5% Sodium deoxycholate and 0.1% SDS) containing protease inhibitors (Roche, Complete Mini EDTA-Free) and phosphatase inhibitors (PhosSTOP, Roche). The lysates were incubated on ice for 20 min and cleared by centrifugation at 4℃ for 15 min at 13,000g. The supernatant was transferred to a new tube for further analysis.

### Immunoprecipitation of Repo-Man in HeLa cells

Prometaphase arrested HeLa cells were resuspended in 200 µL of lysis buffer (50 mM Tris pH 7.5, 150 mM NaCl, 1% NP40, 0.5% Sodium deoxycholate and 0.1% SDS) containing protease inhibitors (Roche, Complete Mini EDTA-Free) and phosphatase inhibitors (PhosSTOP, Roche). The lysates were incubated on ice for 20 min and cleared by centrifugation at 4℃ for 15 min at 13,000 g. The supernatant was then diluted by adding 300 µL of dilution buffer (50 mM Tris pH 7.5, 150 mM NaCl). A total of 500 µL diluted lysates was transferred to a new tube containing pre-washed GFP-Trap magnetic agarose beads (ChromoTek gtma) and incubated at 4℃ for 2 h. After incubation, the beads were washed twice with washing buffer (50 mM Tris pH 7.5, 150 mM NaCl and 0.05% NP40) and once with PBS, then transferred to a clean tube in the last wash. The beads were resuspended in LDS sample buffer (Invitrogen). After heating the beads at 70℃ for 10 min, the eluted proteins were transferred to a new tube for MS analysis.

### Sample preparation for proteomics analysis

All protein samples were sonicated for 10 cycles (30s on/off) using the Bioruptor® Pico, and then centrifugated at 20,000g for 15 min. After adding TCEP (10 mM final concentration, Sigma-Alrich), the supernatant proteins were denatured and reduced at 95℃ for 10 min, and then alkylated with 40 mM IAA (final concentration, Sigma-Alrich) in dark at room temperature for 30 min. The protein concentration was measured using EZQ™ Protein Quantitation Kit (Thermo Fisher Scientific) by following the manual. The proteins from each sample were further processed using SP3 protocol as described ^70^. In brief, protein samples were mixed with SP3 beads (1: 10, protein: beads). For the HILIC fractionation samples from cell lysates, the starting material of proteins was 500 ug. For IP-MS samples, 100 ug of beads were added into the sample regardless of the protein amount. The proteins samples were then incubated and cleaned with SP3 beads, then digested with lysC/trypsin mixture (1: 50, enzyme: protein, Promega) at 37℃ overnight. For IP-MS samples, 50 µL of 2% DMSO (in H_2_O) was used to elute the peptides. The peptide samples were directly used for LC-MS analysis. For the HILIC fractionation samples, after the overnight digestion using lysC/trypsin mixture, PNGase F (1000 U for 1 mg proteins/peptides) was added into the samples for another incubation at 37℃ for 4 h. 200 µL of 0.1% TFA (in H_2_O) was used to elute the peptides from SP3 beads. The peptide concentration was measured using Pierce™ Quantitative Fluorometric Peptide Assay (Thermo Fisher Scientific) by following the manual.

### HILIC fractionation

The 200 µL of peptide samples were added with 200 µL of 0.1% Trifluoroacetic acid (TFA) in Acetonitrile (ACN) and mixed well to have a clear solution. 3 more times of 0.1% TFA (in ACN) were added to reach a final concentration of 80% ACN with 0.1% TFA. It’s crucial to have a clear solution before HILIC fractionation. Centrifugation might be needed to remove possible insoluble content. HILIC was performed using a Dionex RSLCnano HPLC (Thermo Fisher Scientific). Peptides were injected onto a 4.6 mm ID × 15 cm TSKgel Amide-80 column (3 um pore size), using a 90-min multistep gradient with a constant flow rate of 0.4 mL/min: 0-20 min, 80%B; 20-30 min, 80%-70%B, 30-60 min, 70%-60%B, 60-65 min, 60%-0%B, 65-70 min, 0%B (flow rate at 0.2 mL/min). The mobile phases were 0.1% TFA in H_2_O for solvent A and 0.1% TFA in ACN for solvent B. Fractionations were collected by every 126 s, starting from 3 min to 70 min. All the peptide fractions were then dried using SpeedVac concentrator (45℃ or lower temperature, Thermo Fisher Scientific). The samples could be stored in -20℃ or resuspended in 0.1% FA and directly used for LC-MS analysis.

### LC-MS/MS analysis

All the HILIC fractionation and IP samples were analysed by using a Q Exactive Plus mass spectrometer platform (Thermo Fisher Scientific), equipped with a Dionex Ultra-high-pressure liquid-chromatography system (RSLCnano). RPLC was performed using a Dionex RSLCnano HPLC (Thermo Fisher Scientific). Peptides were injected onto a 75 μm × 2 cm PepMap-C18 pre-column and resolved on a 75 μm × 50 cm RP-C18 EASY-Spray temperature-controlled integrated column-emitter (Thermo Fisher Scientific), using a 2-hour multistep gradient with a constant flow rate of 300 nl/min. The mobile phases were H_2_O incorporating 0.1% Formic acid FA (solvent A) and 80% ACN incorporating 0.1% FA (solvent B). The gradient for HILIC enriched samples was: 0-6 min, 1%B; 6-10 min, 1%-8%B; 10-80 min, 8%-28%B; 80-95 min, 28%-38%B. The gradient for IP-MS samples was: 0-6 min, 5%B; 6-91 min, 5%-38%B. The MS data were acquired under the control of Xcalibur software in a data-dependent acquisition mode (DDA) using top N mode. The survey scan was acquired in the orbitrap covering the m/z range from 375 to 1,600 with a mass resolution of 70,000 and an automatic gain control (AGC) target of 3e^6^ ions with 20 ms maximum injection time. For HILIC peptide fractions, the top 20 most intense ions were selected for fragmentation using HCD with 27% NCE collision energy and an isolation window of 1.6 Da with a dynamic exclusion of 60 s. The MS2 scan was acquired in the orbitrap with a mass resolution of 17,500 with the m/z range starts from either 80 or 100. The AGC target was set to 1e5 with a maximum injection time of 60 ms. For IP-MS samples, the target PRM mode was applied for fragmentation, with a AGC target of 2e^5^ and a maximum injection time of 120 ms.

All the synthetic peptides were analysed using an Orbitrap Fusion Tribrid mass spectrometer platform and RSLCnano system. The gradient was: 0-6 min, 5%B; 6-31 min, 5%-38%B; 31-35 min, 38-95%B; 35-40 min, 95%. The MS data were acquired under the control of Xcalibur software in a target PRM. The survey scan was acquired in the orbitrap covering the m/z range from 375 to 1,575 with a mass resolution of 60,000 and an automatic gain control (AGC) target of 4e^5^ ions with 50 ms maximum injection time. The target ions were selected for fragmentation using HCD with 30% NCE collision energy and an isolation window of 1.6 Da with a dynamic exclusion of 60 s. The MS2 scan was acquired in the orbitrap with a mass resolution of 30,000 with the m/z range starts from 80. The AGC target was set to 5e4 with a maximum injection time of 54 ms. The PRM inclusion list was shown in Table S6.

### MS Data analysis

The MS data were analysed together using MaxQuant ^71,72^ (v. 2.3.1.0). The FDR threshold was set to 1% for each of the respective Peptide Spectrum Match (PSM) and Protein levels. The data was searched with the following parameters: quantification type was set to LFQ, stable modification of carbamidomethyl (C), variable modifications, oxidation (M), acetylation (protein N terminus), and hydroxylation (P), with maximum of 2 missed tryptic cleavages threshold. First, proteins and peptides were searched against the homo sapiens database from UniProt (SwissProt April 2023) without adding hydroxylation (P) as a variable modification. Then a fasta file was generated based on the identified protein list and used to identify the hydroxylation (P) peptides and sites. The result tables generated from MaxQuant were processed and filtered using Perseus ^73^ (v. 1.6.15.0). For SILAC data analysis, multiplicity for quantification was set to 2 with maximum 3 labelled AAs. “Arg6” and “Lys4” were selected in the heavy label channel.

### *In vitro* hydroxylation assay and dot blot

For *in vitro* hydroxylation, either 3 µM of Repo-Man synthetic peptide (M602 oxidised) (KPLLSPIPELPEVPEM(OX)TPSIPSIRR), or 3 µM of HIF1α peptide

(LDLEMLAPYIPMDDD), was incubated +/− 300 nM of recombinant PHD1/EGLN2 (Active motif #81064). The hydroxylation reaction was performed in a final volume of 30 µL of 20 mM Tris-HCl pH 7.5, 5 mM KCl, 1.5 mM MgCl_2_, 1 mM DTT, 100 µM α-Ketoglutarate (Sigma #75890), 100 µM L-ascorbic acid (Sigma #A7506) and 50 µM Ammonium iron II sulphate hexahydrate (Fluka #09719), for 2.5 h at 30 °C. After incubation, reactions were stopped by addition of 1 mM EDTA and samples subjected to LC-MS/MS analysis. In the case of HIF1α, 5 µL of the reaction was dot blotted onto a nitrocellulose membrane. Serial dilutions of P564 hydroxylated HIF1α peptide, (LDLEMLAP(Hydro)YIPMDDD), were dot blotted in the same membrane as a positive control. Membrane was air dried and blocked with Intercept Blocking buffer TBS (#92760001 0LiCOR) and incubated overnight with anti-HIF1α OH-P564 antibody (1/1000 dilution). After three washes with TBS_T of 5 min each with the last wash being done with TBS, the membrane was incubated with the secondary antibody donkey anti-rabbit IRDye 680RD (926-68073 and imaged using a Typhoon Fuji Imager.

### siRNA transfections of PHD isoforms

RCC4 cells were transfected with 20 μM siRNA of each PHD isoform using Interferin (Peqlab). A random scrambled sequence was used as the control. The siRNA sequences used were previously presented in ^74^.

### Flow cytometric analysis of cell cycle distributions

Adherent RCC4 cells were harvested, pooled, and washed once in PBS prior to being fixed in ice-cold 70% (v/v) ethanol/distilled water. Afterward, the cells were twice washed in PBS and resuspended in Guava Cell Cycle Reagent (Luminex Corp.; #4500-0220) for 30 minutes at room temperature. The cell cycle distribution with a Guava easyCyte HT machine and software. Cells with DNA content between 2N and 4N were allocated into phases G1-, S-, or G2/M of the cell cycle. The cell profiles were gated according to the control in each independent experiment and the number of cells in each phase are expressed as a percentage of the total number of cells counted.

## Acknowledgements

We thank all our collaborators who worked on the project as well as Alejandro Brenes and Michael Batie for all the insightful discussions. We would also like to thank all members of the Lamond Laboratory. This work was funded by the Wellcome Trust grant (<u>206293/Z/17/Z</u>) with additional support provided by grants from BBSRC (BB/V010948/1; APP3732) & UKRI (EP/Y010655/1).

The mass spectrometry proteomics data have been deposited to the ProteomeXchange Consortium via the PRIDE ^75^ partner repository with the dataset identifier PXD044783 (RCC4) and PXD044663 (HEK293).

---

**Figure S1:**
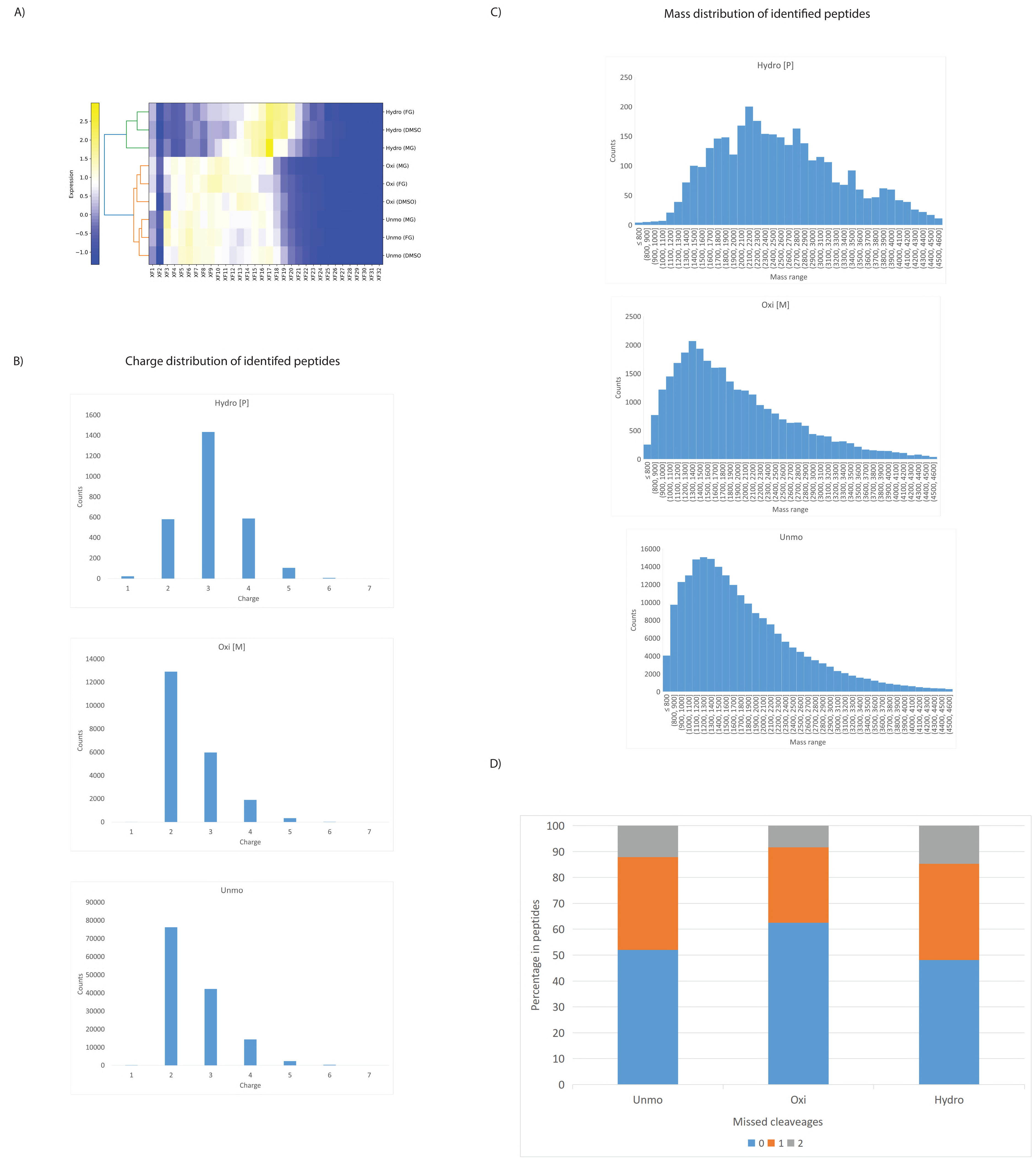
RCC4 dataset, A) The hydrophilicity difference of peptides with hydroxylation (P), oxidation(M) and unmodified peptides across HILIC fractions from all experiments with different treatment. To generate the heatmap, “modificationSpecificpeptides.txt” result file was used and divided into three files for identified peptides with either hydroxylation (P), oxidation(M) or unmodified peptides. The value of “Fraction x” column from of each peptide was used and summarized for each fraction, which represented the total quantity of peptides in each fraction. In each condition, the summarised fraction value of F1, F2, …F24 was scaled to 100% in total. The scaled fraction value were used for heatmap analysis by Heatmapper2 (Nucleic Acids Res. 2025 May 05. doi:10.1093/nar/gkaf385). The hydrophilicity increases from the Fraction 1 to Fraction 32 (XF1…XF32). B) The charge distribution of peptides with hydroxylation (P), oxidation (M) and unmodified peptides. C) The mass distribution of peptides with hydroxylation (P), oxidation (M) and unmodified peptides. D) The missed cleavage difference of peptides with hydroxylation (P), oxidation (M) and unmodified peptides.

**Figure S2:**
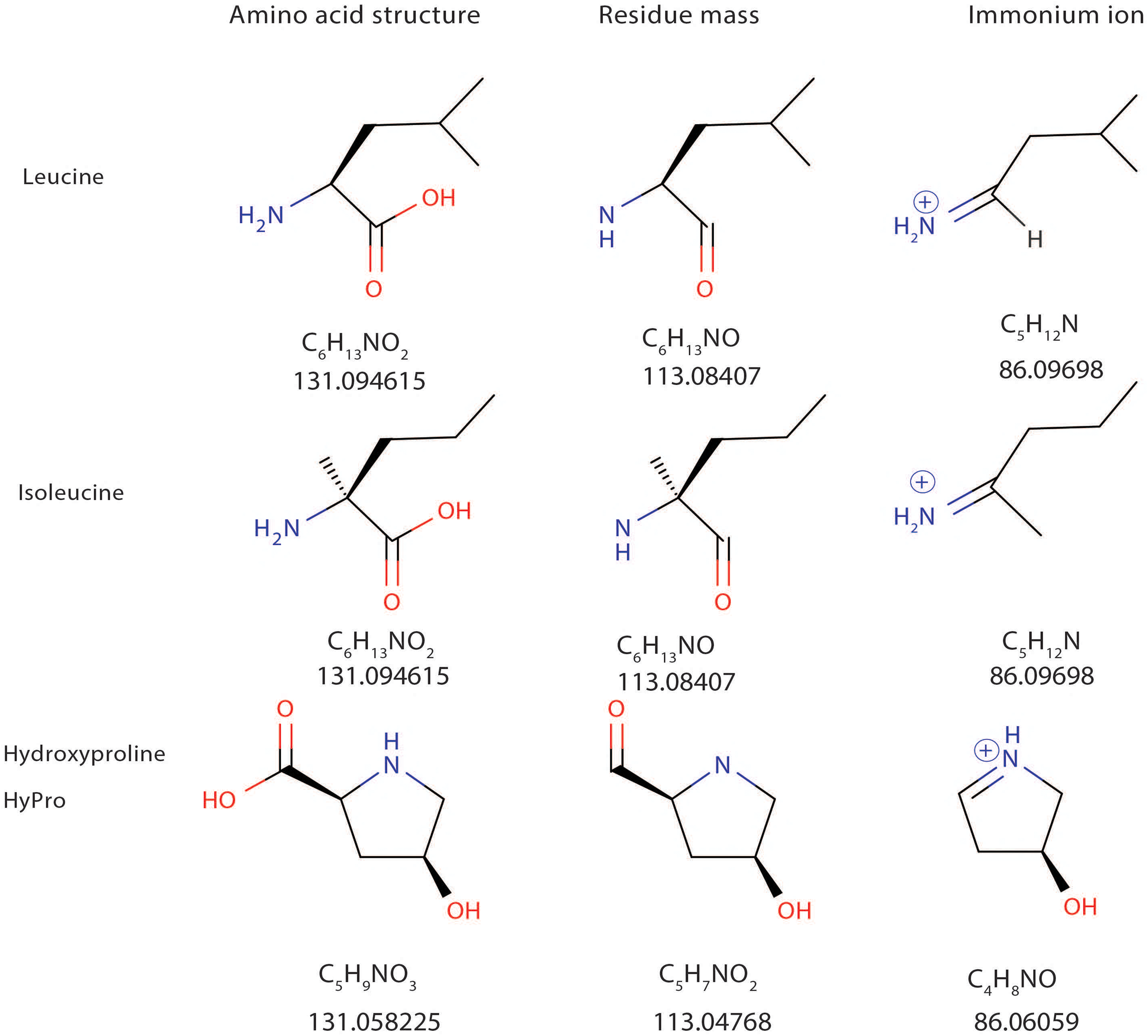
The structure, mass and immonium ion difference of Leucine, isoleucine and hydroxyproline.

**Figure S3:**
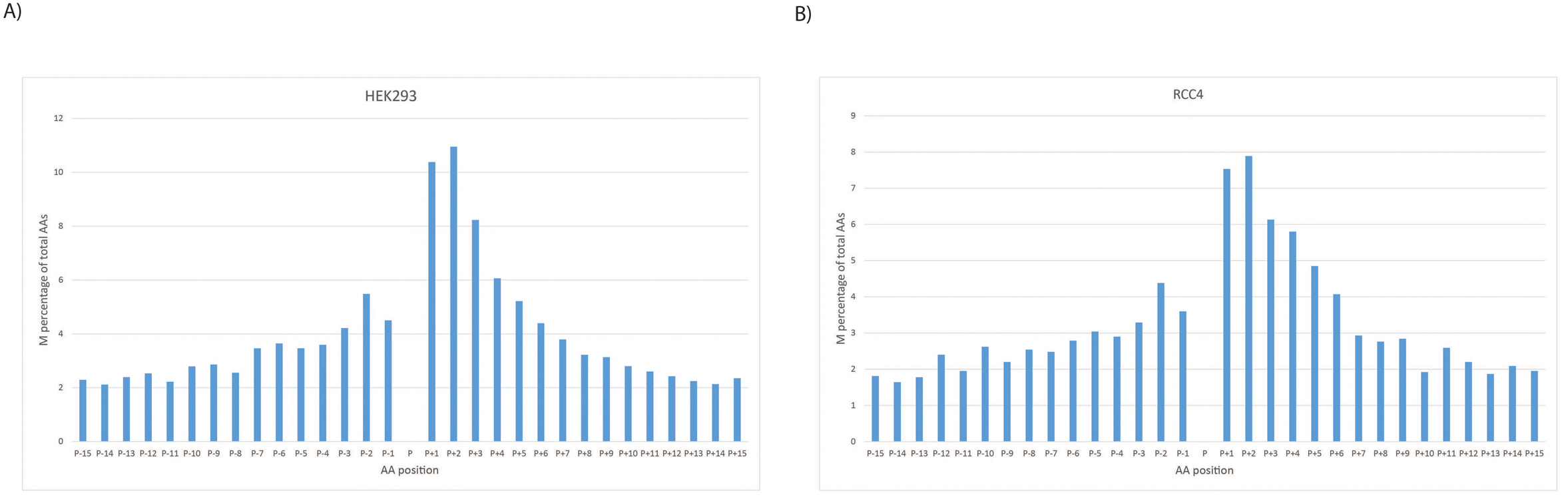
The percentage of M in total amino acids at each position from A) HEK293 and B) RCC4 dataset, centred at the hydroxyproline residue.

**Figure S4:**
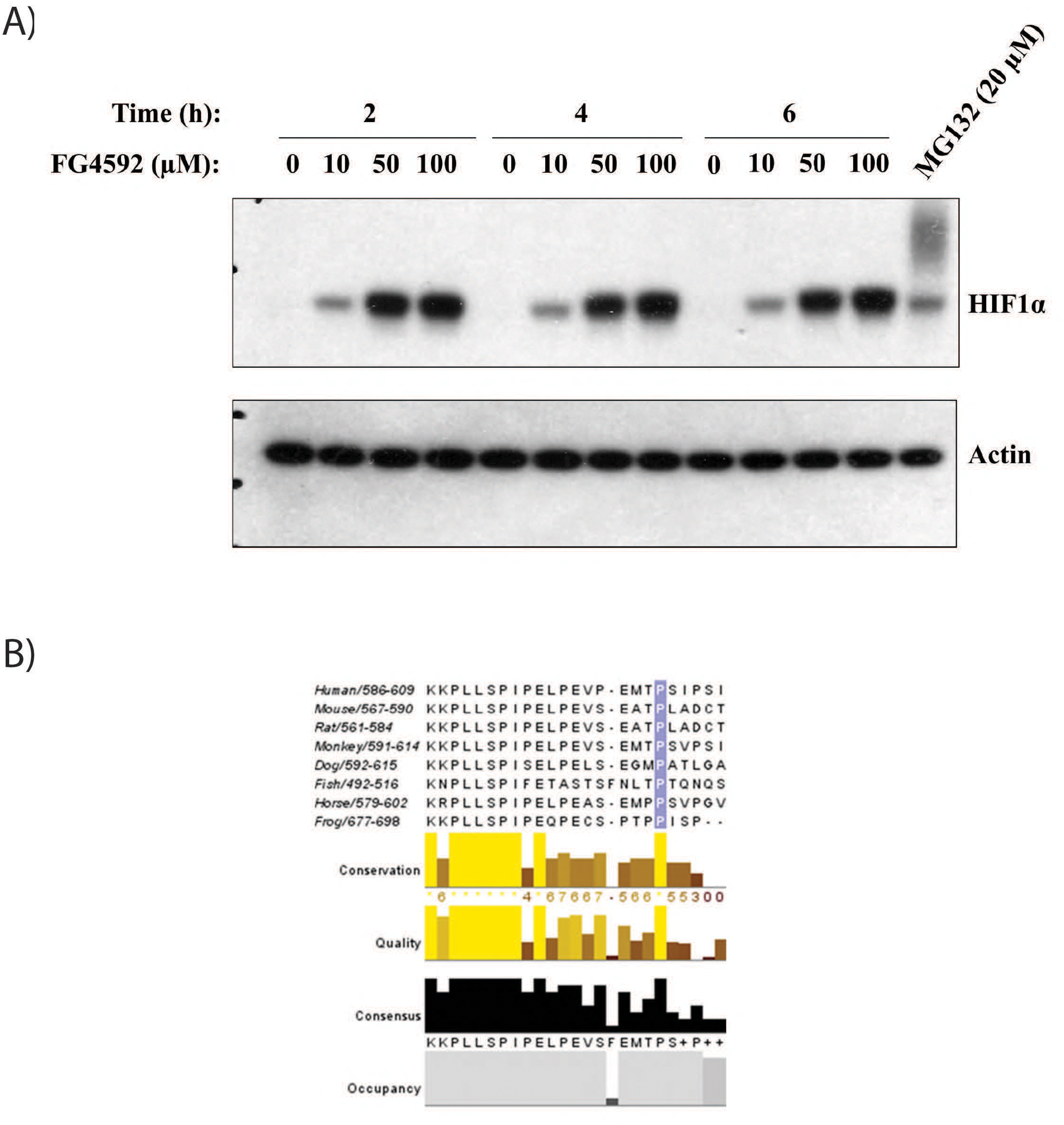
A) Western blot analysis of HIF1α from HEK293 cell lysates treated with different concentrations and times of FG-4592 as indicated in the figure. Actin was used as a loading control. MG-132 was added for 3 h before cells were harvested. B) Sequence alignment of Repo-Man peptide region (covering P604) across different species is displayed.

