## Supplemental materials for "Systematic characterization of site-specific proline hydroxylation using hydrophilic interaction chromatography and mass spectrometry"

Table S1: All hydroxylated proline sites of HEK293 and RCC4 dataset.

Table S2: High confident hydroxylated proline sites of HEK293 and RCC4 dataset. Score >=40 and Localization prob>0.5. Diagnostic peak of HyPro highlighted.

Table S3: High confident hydroxylated proline sites of HEK293 and RCC4 with the removal of M. Score >=40 and Localization prob>0.5.

Table S4: High confident hydroxylated proline sites of HEK293 and RCC4 (FG inhibits), and the overlapping sites from both datasets. Score >=40 and Localization prob>0.5.

Table S5: Reactome pathway results of proline hydroxylated proteins from HEK293 and RCC4 dataset, and the overlapping hydroxylated proteins from both datasets.

Table S6: PRM inclusion list of the target hydroxylated peptides from Repo-man, HIF1α, CEP192 and PKM2.
